# The connectome is necessary but not sufficient for the spread of *alpha*-synuclein pathology in rats

**DOI:** 10.1101/567222

**Authors:** Miguel A.P. Oliveira, Sylvain Arreckx, Donato Di Monte, German A. Preciat, Ayse Ulusoy, Ronan M.T. Fleming

**Affiliations:** Luxembourg Centre for Systems Biomedicine, University of Luxembourg, 6 Avenue du Swing, Belvaux, L-4362, Luxembourg; German Center for Neurodegenerative Diseases, Sigmund-Freud-Strasse 27, D-53127, Bonn, Germany; Analytical Biosciences, Division of Systems Biomedicine and Pharmacology, Leiden Academic Centre for Drug Research, Leiden University, Leiden, The Netherlands

**Keywords:** Parkinson’s disease, animal models, *alpha*-synuclein, connectome, neuronal vulnerability

## Abstract

Certain neuronal populations are selectively vulnerable to *alpha*-synuclein pathology in Parkinson’s Disease, yet the reasons for this selectivity are unclear. Pathology affects neuronal populations that are anatomically connected although the contribution of neuronal connectivity remains to be quantitatively explored. Herein, we simulate the contribution of the connectome alone to the spread of arbitrary aggregates using a computational model of temporal spread within an abstract representation of the mouse mesoscale connectome. Our simulations are compared with the neuron-to-neuron spread of *alpha*-synuclein that has been observed with *in vivo* spreading experiments in rats. We find that neuronal connectivity appears to be compatible with the spreading pattern of *alpha*-synuclein pathology however, it may be *per se* insufficient to determine the anatomical pattern of protein spreading observed in experimental animals, suggesting a role of selective vulnerability of neuronal pathways to *alpha*-synuclein diffusion, accumulation and pathology.

**Graphical abstract:** 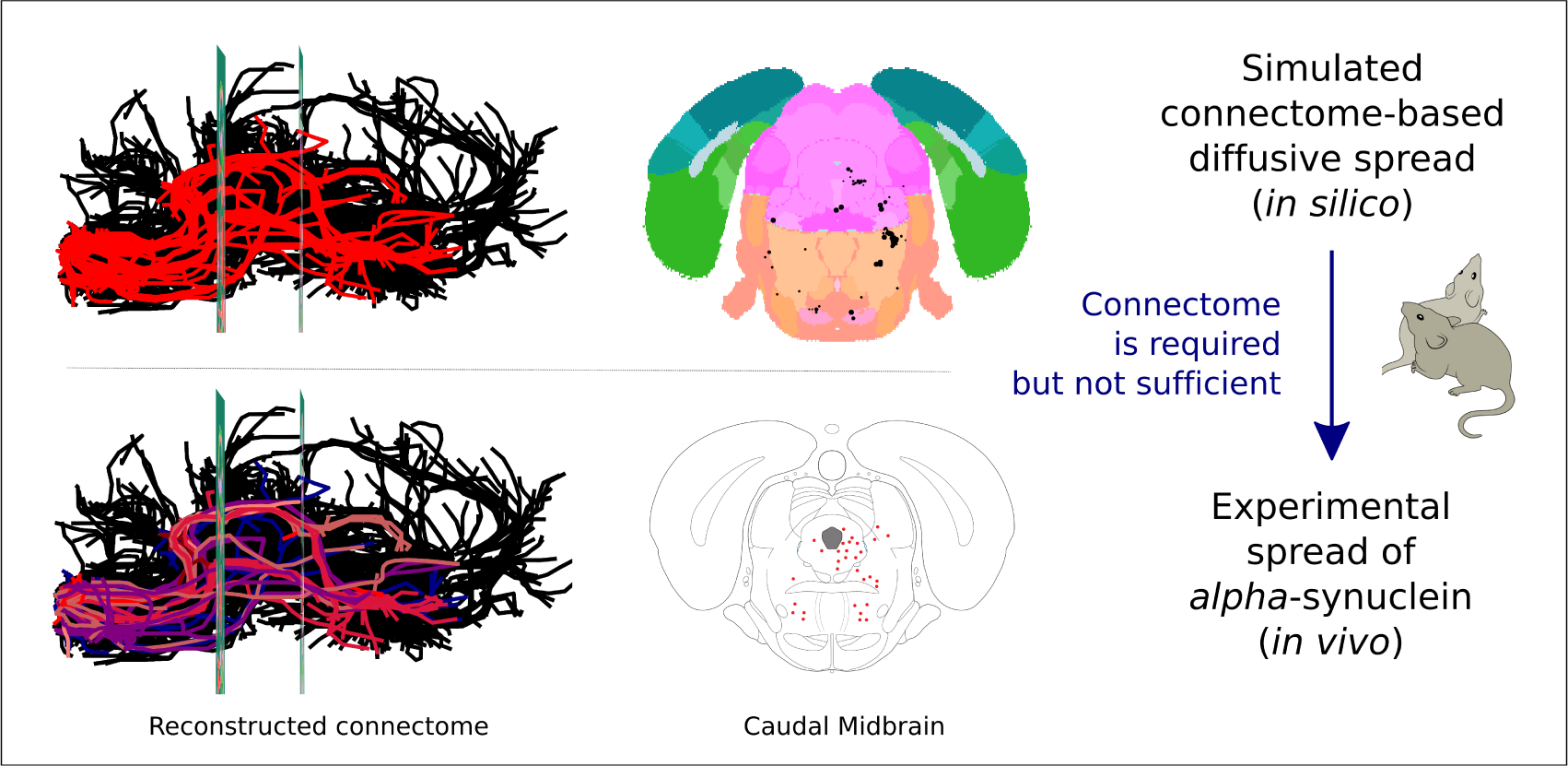

## Introduction

Neurodegenerative diseases progress over time and are generally characterised by specific and clinically identifiable symptoms, such as motor dysfunction, e.g., bradykinesia, or problems of mental function, e.g. dementia. A variety of neurodegenerative diseases have been related to the aggregation of misfolded proteins and transmission of the corresponding proteinaceous seeds (Brundin et al. 2010). *In vitro*, these seeds can corrupt the folding of endogenous proteins and their inter-neuronal transmission could be associated, at least in part, with the progression of pathology (Guo et al. 2014). The spread of pathology from affected to unaffected neuronal populations resembles a “prion-like” sequential dissemination (Brettschneider et al. 2015). This dissemination can occur either locally, between neighbouring cell bodies as a result of exocytosis or cell death, or through the neuronal network, due to synaptic release or uptake (Frost et al. 2010). Examples of disease-specific misfolded proteins are *alpha*-synuclein in Parkinson’s disease (PD), *β*-amyloid and tau in Alzheimer’s disease, huntingtin in Huntington’s disease, and Ataxin-3 in Machado-Joseph disease. In contrast to prion diseases, which are transmitted by an infectious misfolded protein, the aforementioned neurodegenerative diseases seem to originate from *de novo* aggregation of misfolded proteins and spread of pathology through a disease-specific sets of neuronal populations (Jucker et al. 2013).

In many neurodegenerative diseases, pathology primarily affects a disease-specific subset of disperse neuronal populations and results in the accumulation of intra- or extra-neuronal deposits and, ultimately, selective neurodegeneration (Fu et al. 2018). This selective vulnerability is not yet fully understood, although recent evidence suggest that it might result from both neuronal connectivity and intrinsic characteristics of the connected neurons, such as morphological characteristics and biochemical profiles (Brettschneider et al. 2015). This hypothesis is of particular interest in Parkinson’s disease (Surmeier et al. 2017; Borghammer 2018), which is defined clinically by motor (e.g., bradykinesia, resting tremor and rigidity) and non-motor (e.g., sleep disorders, olfactory deficits and depression) symptoms (Postuma et al. 2015). Histopathologically, Parkinson’s disease is characterised by intraneuronal accumulation of *alpha*-synuclein-containing inclusion and degeneration of specific neuronal populations, such as dopaminergic cells in the substantia nigra pars compacta (SNc) (Dickson et al. 2009). The onset of Parkin-son’s disease is hypothesised to be up to 20 years before the occurrence of motor symptoms (Hawkes et al. 2010) and the existence of this prodromal phase is supported epidemiologically (Hawkes 2008), by clinical observation of early non-motor symptoms (Goetz et al. 2008), and by a progressive appearance of extra-nigral intracellular protein aggregates that are generally co-identified with *alpha*-synuclein and termed Lewy pathology (Fahn et al. 2004).

Based on the distribution of Lewy pathology in the brain (and other tissues) (Hawkes et al. 2010) a neuropathological temporal staging scheme has been proposed for Parkinson’s disease progression (Braak, Tredici, et al. 2003; Braak, Ghebremedhin, et al. 2004; Braak, Bohl, et al. 2006; Braak and Tredici 2008). Despite variation in the association between Lewy pathology and onset of clinical signs, often non-motor symptoms appear before motor symptoms in a manner consistent with Braak’s neuropathological staging scheme (Braak and Del Tredici 2009). This anatomically specific and consistent picture of protein aggregate progression provides evidence that certain neuronal populations are selectively vulnerable to *alpha*-synuclein pathology in Parkinson’s disease (Halliday et al. 1990; Del Tredici et al. 2002; Sulzer et al. 2013) and also suggests that the progression of the disease may follow a disease-specific pattern that resembles brain connectivity. This pattern further suggests that the disease is transmitted along neuronal pathways and highlights the importance of studying the spread of pathology with respect to neuronal connectivity.

The connectome is a comprehensive map of neural connections within an organism’s nervous system. The structural or functional study of connectomes may range in scale from a full set of morphologically resolved neurons within a part of the nervous system to a macroscopically resolved description of the fibre tracts between different parts of the nervous system. In humans connectomes can be inferred from various methods, including resting-state and task-based functional magnetic resonance imaging (Van Dijk et al. 2010) and diffusion magnetic resonance imaging (Griffa et al. 2013). However, these techniques do not yet admit resolution of individual neuronal projections. Such *mesoscale resolution* can only currently be achieved with post-mortem brain tissue sections (Van Dijk et al. 2010; Castellanos et al. 2013; Fornito et al. 2016; Ghosh et al. 2013; Griffa et al. 2013; Jbabdi et al. 2013; Morgan et al. 2013). Nevertheless, using an inferred human brain connectome, Raj et al. (Raj, Kuceyeski, et al. 2012) developed a network diffusion model of disease progression in Alzheimer’s disease, and used it to investigate the role of brain structure on disease progression (Raj, LoCastro, et al. 2015). Further efforts in this direction, with connectome models of higher resolution, are necessary to investigate the spread of cerebral pathology in neurodegenerative diseases.

Neuropathological evidence of neuron-to-neuron *alpha*-synuclein transmission (Masuda-Suzukake et al. 2013) has been confirmed in multiple mouse and rat spreading models (Ulusoy, Rusconi, et al. 2013; Helwig et al. 2016; Rey et al. 2016; Mason et al. 2016). In such *in vivo* spreading models, *alpha*-synuclein pathology is induced by injection of *alpha*-synuclein fibrils or viral vectors over-expressing exogenous human *alpha*-synuclein into specific brain structures, and spread is inferred by histopathological comparison of cerebral sections from animals sacrificed at different durations after injection. Importantly, the spread of *alpha*-synuclein has been observed following induction of *alpha*-synuclein accumulation in brain structures that are known to have Lewy pathology in early Parkinson’s disease, such as the olfactory bulb (Rey et al. 2016; Mason et al. 2016) and the dorsal motor nucleus of the vagus nerve (DMX) (Helwig et al. 2016; Ulusoy, Rusconi, et al. 2013), which may mimic early PD (Braak, Del Tredici, et al. 2002). These *in vivo* spreading experiments have also been performed by injecting *alpha*-synuclein fibrils into other brain structures involved in Parkinson’s disease, such as the striatum, cortex and substantia nigra (Luk, V. Kehm, et al. 2012; Masuda-Suzukake et al. 2013; Luk, V. M. Kehm, et al. 2012).

To complement these qualitative analyses, a quantitative evaluation of the contribution of neuronal network connectivity to the pattern of spread is required. A comprehensive mesoscale reconstruction of the mouse neuronal connectivity was generated by the Allen Institute connectivity project (Oh et al. 2014), which mapped axonal projections emanating from more than 590 anatomically distinct brain regions using intracerebral injections of a pan-neuronal adeno-associated viral vector expressing enhanced green fluorescent protein. This computational resource includes neuronal projections that cover almost the entire mouse brain, with enough sensitivity to detect individual thin axon fibres.

Herein, we quantitatively predict the contribution of the cerebral connectome to the spread of *alpha*-synuclein pathology using a computational model of spatiotemporal spread along a reconstructed mouse mesoscale connectome (Oh et al. 2014). Furthermore, we compare simulated spatiotemporal spread from vagal nerve connected brain structures, in particular the DMX, with the spread of *alpha*-synuclein pathology observed in an *in vivo* caudo-rostral spreading model, where neuron-to-neuron spreading of exogenous human *alpha*-synuclein is initiated by injecting viral vectors into the vagus nerve (Helwig et al. 2016; Rey et al. 2016; Mason et al. 2016; Luk, V. Kehm, et al. 2012; Ulusoy, Rusconi, et al. 2013). We address a selection of connectivity related questions. For example, is neuronal network topology sufficient, or necessary, or both, to determine whether a neuronal population is vulnerable to *alpha*-synuclein pathology within *in vivo* models? Does the number or length of neuronal projections between a pair of brain structures determine the propensity for spread of *alpha*-synuclein pathology?

## Materials and methods

An overview of the methodology is given in Figure 1, and further described below.

**Figure 1:**
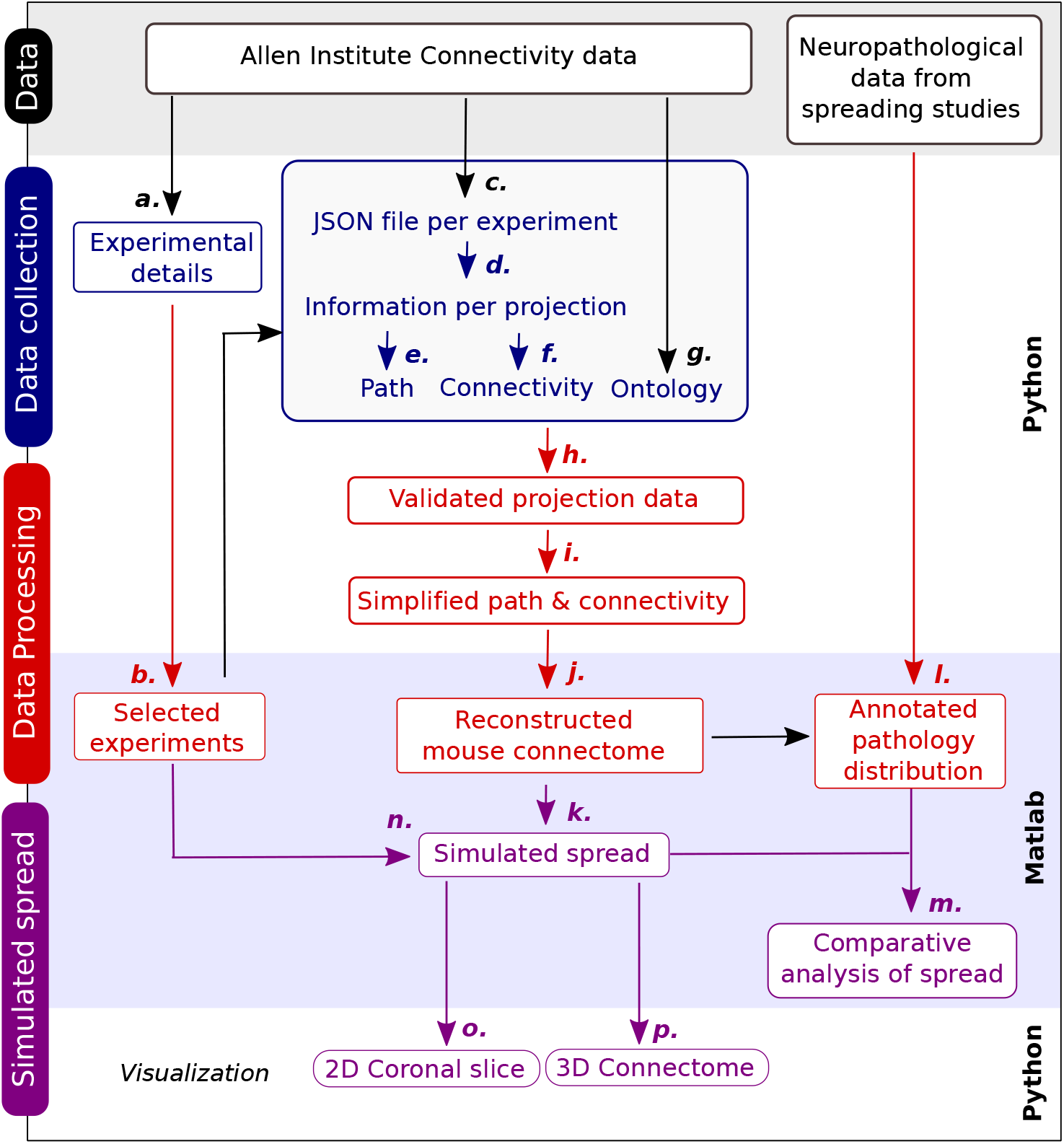
Methodological overview. We reconstructed a whole-brain composite mouse mesoscale connectome, from published data (Oh et al. 2014), then converted it into an *in silico* model that simulates the spatiotemporal diffusion of *alpha*-synuclein *in vivo*. The *in silico* model predictions were compared with neurohistopathologically evaluated exognous *alpha*-synuclein staining in rats that were previously injected in the vagal nerve with an rAAV virus encoding for human *alpha*-synuclein. Cerebral neuronal connectivity data was obtained from the Allen Brain Institute Connectivity project. Metadata on all connectivity experiments (a) was unified and analysed to select appropriate experiments to build the reconstruction (b). Data on each connectivity experiment, was obtained from one JSON file (c), and contains information on neuronal projections to multiple target structures, from one, or often multiple, injected source structures (d). For each neuronal projection, its path coordinates (e) and the coordinates for the corresponding source and target structures were collected (f). The Allen Brain Institute *software developmental kit* (SDK) was used to obtain a correspondence between brain structure coordinates and ontology (g). For each experiment, each pair of uniquely connected structures was identified, invalid projections were discarded and the full path coordinates for each valid projection were interpolated (h). For each pair of uniquely connected structures, one model projection was selected (i) then model projections from all selected experiments were then assembled to reconstruct an anatomically localised, composite, mesoscale, mouse connectome (j). The spread of exogenous *alpha*-synuclein was simulated as a spatiotemporal diffusion process along the connectome (k). Quantitatively predicted *alpha*-synuclein accumulation was compared with semi-quantitative data from independent *alpha*-synuclein spreading experiments in rats (l), for each brain structure independently (m) and for *in silico* and *in vivo* coronal slices of the mouse brain (o). Simulated spread was also visualised in three dimensions for comparison with future experimental data (p). Due to the high density of model projections, when visualising images or animations of three dimensional simulated spread for the whole brain, we selected a subset of connectivity experiments that were initiation structures in spreading experiments, either as source or target structures (n).

### Reconstruction of a whole-brain composite mesoscale connectome

The Allen Reference Atlas structures form a contiguous spatial subdivision of the brain, and each encompasses a three dimensional volume, delineated by an anatomically determined boundary. Each Allen Brain Atlas connectivity experiment returns data on a multitude of neuronal projections (Supplementary Section 1.1 and 1.2). Each neuronal projection has an origin within a source focus and a terminus in a target brain structure. When a source focus also included one, or two, secondary source structures, we assumed that ½, or ⅓, of the neuronal projections initiated in each structure, respectively. When a source structure was a parent in the ontology, the number of projections were further subdivided equally among the corresponding leaf structures. In this manner, we resolved all projections to the level of leaf source and target structures.

In order to simulated spread across a whole-brain, in a computationally tractable way, the data from each connectivity experiment was simplified in a standardised manner. A pair of neuronal projections are said to be *similar*, if they arise from the same connectivity experiment, and they connect the same pair of leaf brain structures. In each experiment, between any pair of brain structures, a single *model projection* was used for modelling and visualisation, and all similar neuronal projections were discarded (Figure 1i). A model projection was defined as the one with the length closest to the average length over all similar projections. Nevertheless, the actual number of similar projections was used as a parameter for each model projection in the *in silico* spreading model (*w* ∈ ℝ^*m*^ in 2 below). Therefore, we represent the mouse brain connectome by representative neuronal projections, between pairs of representative brain structures.

Each model projection follows a directed trajectory from a *source path vertex* within a source brain structure, through a sequence of non-equidistant intermediate points, to a *target path vertex* within a target brain structure. We used B-spline interpolation, followed by linear segmentation, into 50*μm* segments, to decompose each model projection into a sequence of linear, directed *path edges* connecting a sequence of *path vertices* (Figure 1i).

Each brain structure is represented by one *structure vertex* localised to one central point within that mouse brain structure. To connect each model projection to a pair of model structures, a directed *structure edge* extends linearly from the *source structure vertex* to the *source path vertex* of a model projection, and another directed *structure edge* extends linearly from the target path vertex of the same model projection to a target structure vertex. These structure edges are modelling constructs, necessary to ensure connectivity between pairs of model projections, due to saturation of the fluorescence signal within the source focus, which hinders resolution of individual projections.

The mesoscale mouse brain connectome (Figure 1j) is represented by a *connectome graph* 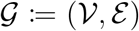 between a set of vertices, or nodes, denoted 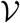 and a set of edges, or links, denoted *ℰ*. There are 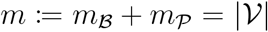 vertices, where 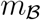 is the number of structure vertices and 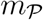 is the total number of path vertices for all projections. Path vertices are labelled *P_i_* and brain structure vertices are labelled as *B_i_* (see Figure 11a). There are 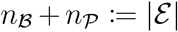 edges, where 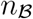 is the number of structure edges and 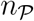 is the total number of path edges for all projections. Each edge is defined with an anterograde orientation, but simulation of the spread of *alpha*-synyclein, described next, is invariant with respect to this conventional assignment.

### Simulating the spread of *alpha*-synyclein

We simulate the spread of *alpha*-synyclein as a discrete time Markov process (Figure 1k). That is, the amount of spread at a given instant is only a function of the amount of *alpha*-synyclein present in that instant, and not before it. Let *N* be the incidence matrix for the connectome graph 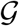. It has 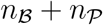 rows, corresponding to vertices, and 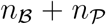 columns, corresponding to edges. The entry in *i^th^* row and *j^th^* column is defined as follows: *N_i,j_* = −1 if 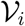 is the source vertex, *N_i,j_* = 1 if 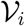 is the target vertex, and *N_i,j_* = 0 otherwise. As each edge is between a pair of vertices, each column of *N* must have exactly two non-zero entries. Let *w* ∈ ℝ^*m*^ denote the weight of each path edge, given by the number of similar projections represented by the corresponding model projection. Let *c*(*k*) ∈ ℝ^*m*^ denote the amount of *alpha*-synyclein at each vertex at time step *k* ≥ 0. A discrete time Markov process starts in an initial state *c*_0_, and steps to the next state based on

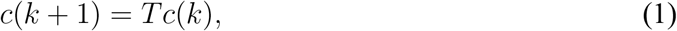

where

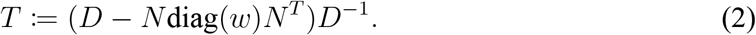

The entries *T_i,j_* are referred to as transition probabilities, and the matrix is a transition matrix *T*. Additionally, *D*:= diag(*d*), with *d* ∈ ℕ^*m*^ where *d_i_* is the degree of vertex 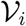, that is *d* = diag(*w*)abs(*N^T^*)**1**. Equation 1 is applied successively to predict the diffusive spread of *alpha*-synyclein at each time step. Figure 11 illustrates the evolution of simulated spread from time step *k*_0_ to time step *k*_6_ on a small representative network.

### Reconstruction of a first-order composite mesoscale connectome

An established rat model of caudo-rostral *alpha*-synuclein spreading was used for comparison with computational model predictions (Supplementary Section 2). In the rat spreading experiments, the adeno-associated virus, expressing human wild-type *alpha*-synuclein, only infects the neurons that extend projections to the point of injection within the vagal nerve. This includes soma of neurons in cerebral structures (DMX and AMB) that extend projections into the vagus, and projections from vagal ganglionic neurons that project to specific cerebral structures (NTS, SPVC and AP). In this model, *alpha*-synuclein expressed by the adeno-associated virus is found only at the recipient neuron that are directly connected with the initiation structure. Patient data suggest that *alpha*-synuclein is transmitted predominantly in a retrograde manner (Surmeier et al. 2017), however *in vitro* studies have shown that *alpha*-synuclein is capable of being transported in both anterograde and retrograde direction after being taken up either from the axon terminal or soma (Brahic et al. 2016). Therefore we defined a first-order composite connectome composed of all model projections in the whole-brain composite connectome with either a source or target structure in the set of initiation structures (DMX, AMB). When comparing simulated and actual spread in rats, the computational model was set to only allow spread through the first-order composite connectome.

### Comparison of simulated and *in vivo* spread in rat coronal slices

In order to directly compare simulated spread with spatial patterns of *alpha*-synuclein pathology, virtual coronal slices were generated at a set of rostrocaudal positions, corresponding to those selected for histopathological evaluation of *alpha*-synuclein pathology in rat brains (Figure 1o). Each model projection is represented by a sequence of vertices, connected by linear segments of uniform length, with coordinates specified with reference to the Allen Mouse Brain Reference Atlas. Based on the histopathological features of the rat brain slices, the corresponding coronal plane was chosen in the Allen Mouse Brain Reference Atlas. The intersection between a linear segment of a model projection and the coronal plane defines the potential position of virtual *alpha*-synuclein accumulation. Simulated spread was then be displayed, or not, in a virtual coronal section based on the severity (*c_i_*(*k*) > 0), or absence (*c_i_*(*k*) = 0), of simulated *alpha*-synuclein accumulation at time step *k*, for the vertex 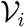 that was the shortest linear distance from the coronal plane in question. In coronal slices, brain structure colours are based on the Allen Brain Atlas ontology, and coronal slices were obtained from the Allen Brain Institute volumetric data compressed into a NRRD (Nearly Raw Raster Data) format.

### Three dimensional whole-brain connectome spread visualisation

Three dimensional visualisations of the simulated spread of *alpha*-synuclein were also generated (Figure 1p). However, due to the high density of model projections, it was difficult to discern patterns or individual model projections when visualising three dimensional animations, or snapshots thereof, for the whole-brain. Therefore, three dimensional visualisations of the evolution of virtual *alpha*-synuclein spread are only presented for projections in the first-order connectome plus the additional projections from connectivity experiments that sent a projection to at least one of the initiation structures (DMX, AMB), in the rat spreading experiments (Figure 2b). Three dimensional animations and snapshots were generated using the Python package `Matplotlib`.

**Figure 2:**
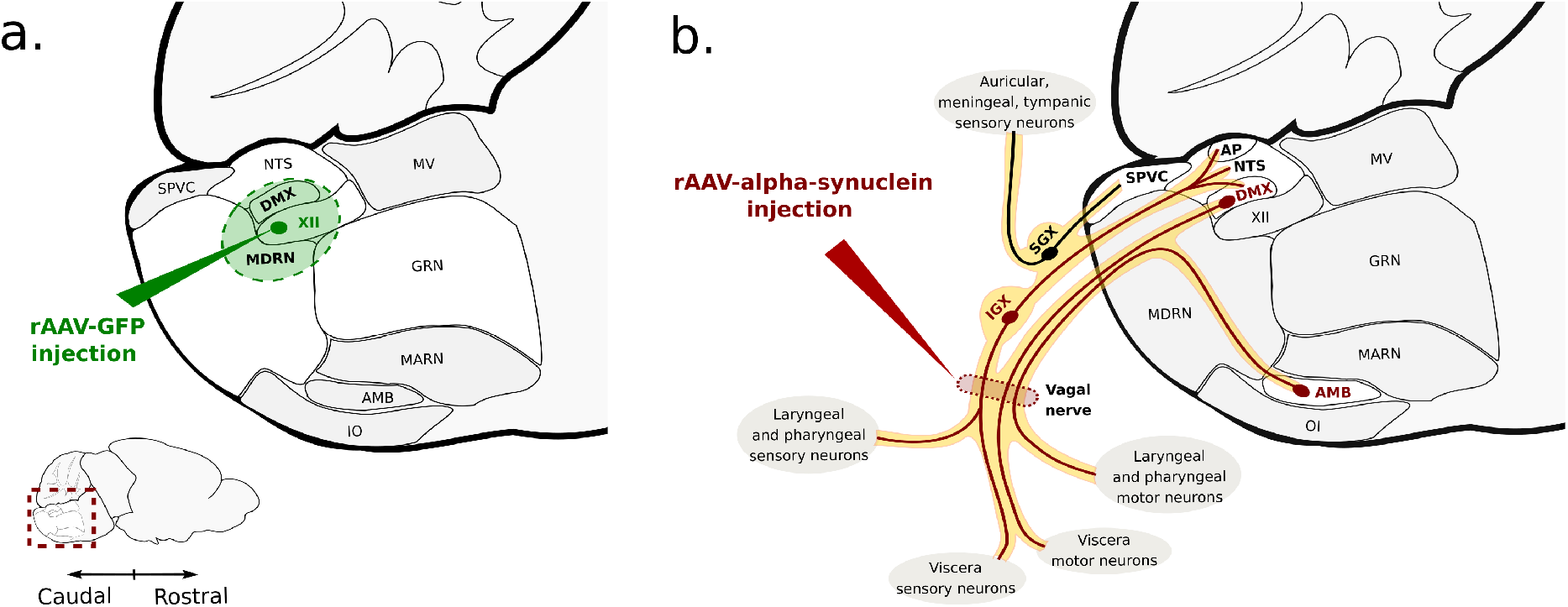
Overview of mouse connectivity and rat spreading experiments. (a) In each published mouse connectivity experiment from Allen Brain Institute (Oh et al. 2014), an injection of an anterograde tracer, recombinant adeno-associated virus (rAAV) expressing enhanced green fluorescent protein (GFP, green arrow), results in a focus of cerebral fluorescence (shaded green), that is maximal in a primary source structure (XII). However, adjacent structures may also fluoresce, two of which, with the most pixels fluorescing, relative to their volume, we designate as secondary source structures (DMX, MDRN). Our composite mouse mesoscale connectome was reconstructed from 207 independent connectivity experiments, with projections emanating from one primary and two secondary structures per experiment (three source structures). (b) In the published rat spreading experiment (Ulusoy, Rusconi, et al. 2013), the left vagus nerve (yellow) is injected with adeno-associated virus, expressing human *alpha*-synuclein, at a position (red arrow) distal to the inferior ganglion of the vagus (IGX). This primarily infects nerve fascicles and tracts (red curves) and neuronal soma (red ovals) of brain structures (DMX, AMB) and ganglionic neurons (IGX) located in the vagal trunk, outside of the central nervous system. Thereby, axons within brain structures that receive vagal projections (NTS, AP, DMX) may also express human *alpha*-synuclein. However ganglionic neurons and their central projections that branch out from the vagal trunk at positions proximal to the injection site, are not directly infected, e.g. the superior ganglion of the vagus (SGX) and the corresponding caudal part of the spinal nucleus of the trigeminal nerve (SPVC).

## Data availability

Connectivity data is available in the Allen Brain Institute (http://connectivity.brain-map.org/) connectivity project (Oh et al. 2014) using its application programming interface (http://help.brain-map.org/display/mouseconnectivity/API). The first report, with a summary of the data from the rat spreading model is available in Ulusoy et al. 2013 (Ulusoy, Rusconi, et al. 2013).

## Results

We generated an *in silico* model to simulate the diffusive spread of virtual *alpha*-synuclein through a composite mesoscale mouse connectome, as a function of both connectome topology and geometry. We simulated spread through a first-order composite mesoscale mouse connectome and compared it to the brain distribution of *alpha*-synuclein pathology in an established *alpha*-synuclein spreading model in rats. Furthermore, we simulated spread of *alpha*-synuclein through a whole-brain composite mesoscale mouse connectome, with initiation in the dorsal motor nucleus of the vagus nerve, and compared it with the reported distribution of Lewy pathology within the brains of Parkinson’s disease patients at early Braak stages.

### Reconstruction of a composite mesoscale mouse connectome

A composite, mesoscale, connectome was reconstructed for a typical male `C57BL/6J` mouse, using neuronal projections identified with a fluorescent anterograde tracer (rAVV-GFP) expressed by a recombinant adeno-associated virus (Oh et al. 2014). This reconstructed connectome used a comprehensive set of 207 independent connectivity experiments, with projections emanating from a set of 439 unique *leaf brain structures*. Here, leaf refers to the highest level of anatomical resolution in the ontological tree used by the Allen Mouse Brain Atlas. In each connectivity experiment, brain structures were considered subject to certain parameters that eliminated projections from any insufficiently represented brain structure. Figure 2a illustrates an example of one of the 207 connectivity experiments (`ABAID:127350480`) where the hypoglossal nucleus (XII) was designated the *primary source structure*, and two adjacent structures, the dorsal motor nucleus of the vagus (DMX) and medullary reticular nucleus (MDRN), were designated *secondary source structures*, based on thresholds on the magnitude and fraction of fluorescently labelled brain structure volume. The contribution of other partially fluorescent brain structures, e.g. the nucleus of the solitary tract (NTS), were omitted from this connectivity experiment. From all 207 connectivity experiments, 307 structures were primary in at least one experiment, while 132 were always secondary structures (Supplementary Figure 7).

After quality control for spurious projections, an average connectivity experiment contained 21, 338 individual neuronal projections, but in average it represents connections between only 1, 277 unique pairs of brain structures (Supplementary Figure 8). Composition of similar projections, sharing the same pair of brain structures in one experiment (Figure 3), enabled the simplification of a total of 3, 169, 406 neuronal projections into 163, 236 model projections between 829 unique leaf brain structures. The Allen Mouse Brain Reference Atlas defines a total of 836 leaf structures within the grey matter and ventricular systems. Therefore, our composite mesoscale mouse connectome represents projections emanating from about half (439/836) of all brain structures to almost all brain structures (829/836) in the actual mesoscale mouse connectome (cf Supplementary Figure 9), yet results in a computational model that is amenable to efficient simulation of diffusive spread of *alpha*-synuclein as a discrete time Markov process.

**Figure 3:**
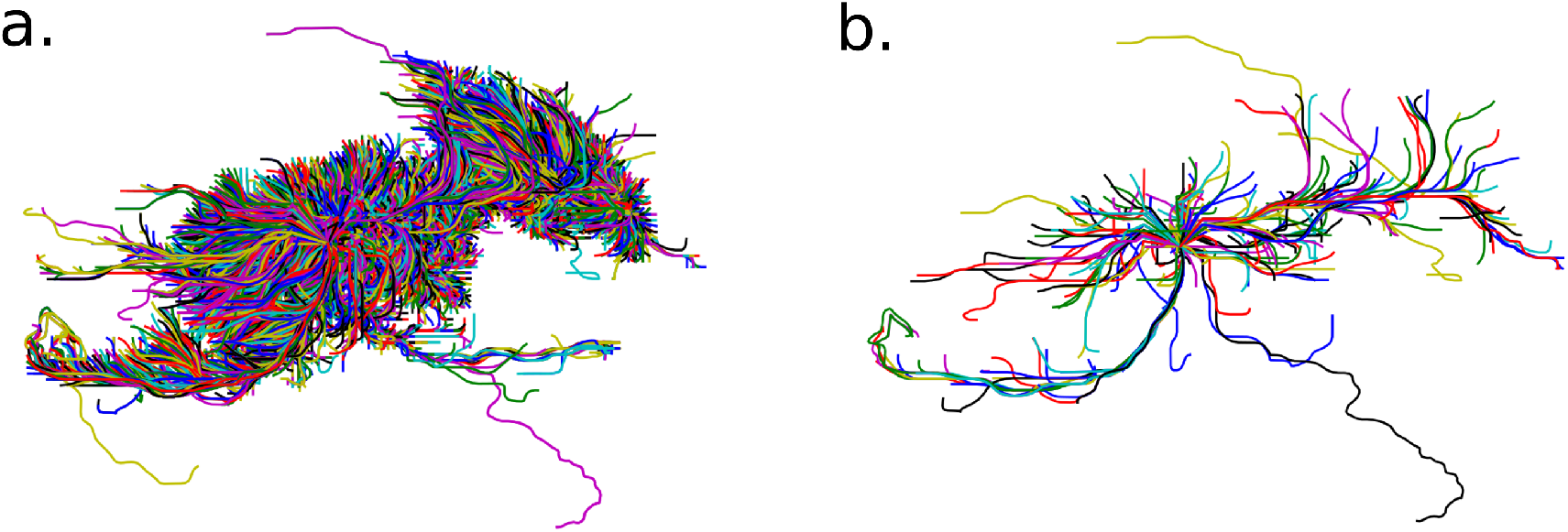
Composition of similar neuronal projections. An example connectivity experiment (ABAID:158914182) with injection site in the reticular part of the substantia nigra (SNr). (a) Before simplification there were 33,600 neuronal projections between 250 unique pairs of brain structures. (b) After composition of similar neuronal projections, there result is one model projection between each of the 250 unique pairs of brain structures.

### Distribution of *alpha*-synuclein pathology in a rat model

The composite, mesoscale, mouse connectome contains 829 leaf brain structures, each of which corresponds to one of 139 rat brain structures that were microscopically resolved in our analysis of histological sections. Within this whole-brain connectome, the first-order connectome with respect to projections to, or from, the initiation structures (DMX and nucleus ambiguus, AMB), contained 867 model projections between 323 unique leaf brain structures, as illustrated in Supplementary Figure 9b. These first-order leaf structures correspond to 90 of the 139 microscopically resolved rat brain structures.Outside the first-order connectome, there are 506 leaf structures in the mouse brain, and they correspond to 49 microscopically resolved rat cerebral structures. A semiquantitative review of coronal sections of rat cerebrum, at 12 weeks after injection (Ulusoy, Rusconi, et al. 2013), revealed *alpha*-synuclein pathology, as detected by exogenous *alpha*-synuclein immunoreactivity above a visual threshold of detection, in the soma of the initiation structures (DMX and AMB) and within neurites in 30 other recipient brain structures (Figure 4). These neurites appeared as fine treads, often containing enlarged bead-like varicosities. Previous studies using the same model showed that presence of *alpha*-synuclein in these neurites leads to time-dependent enlargement of the varicosity volume and long-term pathological consequences in the affected brain regions, including neuronal loss and microglial activation (Ulusoy, Rusconi, et al. 2013; Rusconi et al. 2018).

**Figure 4:**
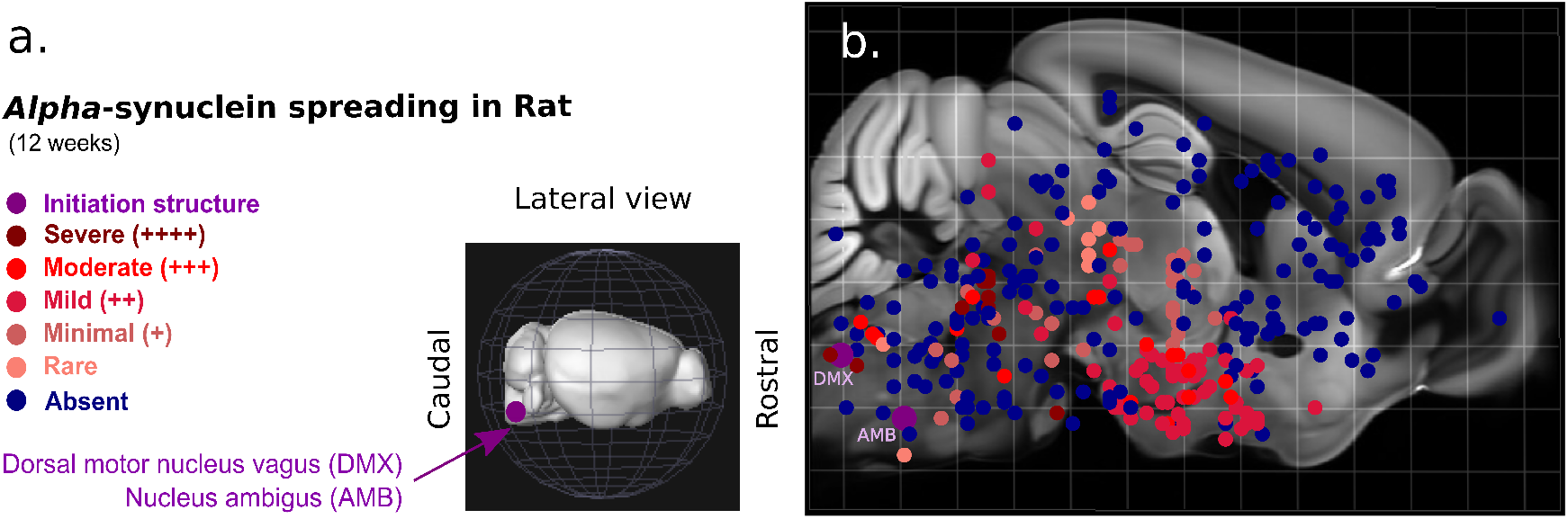
Rat *alpha*-synuclein spreading mapped to a mouse cerebrum. (a) Lateral view of a three dimensional mouse brain with the location of the experimental initiation structures highlighted (purple) (b) Anatomically localised spreading severity, at 12 weeks post-AAV-injection, mapped to homologous leaf structures in the Allen Mouse Brain Atlas (lateral projection onto a mid sagittal section). The initiation structures (DMX and AMB) as well as semiquantitative spreading severity, or absence thereof (blue), are also displayed. See Supplementary Table 1 for details.

All of the microscopically resolved rat brain structures containing *alpha*-synuclein pathology, except the area postrema (AP), correspond to one of the leaf structures in the first-order mouse connectome. That is, the 32 microscopically resolved rat structures with *alpha*-synuclein accumulation correspond to 140 leaf structures in the mouse connectome, and all of them, except the AP, were also in the first-order connectome (Supplementary Table 1). Of the 32 microscopically resolved rat structures with detectable *alpha*-synuclein pathology, 2 displayed severe pathology, the nucleus of the solitary tract (NTS) and the parabrachial nucleus (PB), 6 displayed moderate pathology, e.g., the principal sensory nucleus of the trigeminal (PSV), 10 displayed mild pathology, e.g., the midbrain reticular nucleus (MRN), 8 displayed minimal pathology, e.g., gigantocellular reticular nucleus (GRN), 3 rarely displayed *alpha*-synuclein pathology, e.g., the hypoglossal nucleus (XII).

### Simulated spread of *alpha*-synuclein

Simulation of the spread of *alpha*-synuclein provides a step-wise, anatomically resolved, prediction of virtual *alpha*-synuclein accumulation that is consistent with specified modelling assumptions. Subsequent detailed comparison with histopathological data on *alpha*-synuclein pathology was then used to evaluate the appropriateness of these assumptions. For all simulations of the diffusive spread of *alpha*-synuclein, we assumed that the direction of spread is invariant with respect to the orientation of a neuronal projection and that the rate of spread is proportional to model projection length. That is, the spread of *alpha*-synuclein proceeds along the connectome, either anterogradely or retrogradely, at a rate of one model projection edge per time-step (cf. Figure 11). Also, the fractional spread from one brain structure to another is assumed to be proportional to the amount of projections between them (*w_j_* in Equation 2). That is, all else being equal, the amount of *alpha*-synuclein spread is greater between two brain structures with many individual projections between them, than it would be if there were few.

### First-order simulated spread compared with rat histopathology

We simulated the diffusive spread of *alpha*-synuclein, along the first-order mouse connectome, starting simultaneously from two initiation structures, DMX and AMB. Figure 5 provides a snapshot at the final timestep of the simulation. A three dimensional animation (Supplementary Video hyperlink) illustrates that the evolution of virtual *alpha*-synuclein accumulation can be followed with high spatiotemporal resolution, but for quantitative comparison with cerebral *alpha*-synuclein pathology in the rat spreading model, we focus our comparison on brain structures that were also evaluated histopathologically in rat coronal brain slices. Here we assume that virtual *alpha*-synuclein accumulation in a coronal slice can be attributed to a brain structure if virtual *alpha*-synuclein reaches the corresponding source or target structure, in that virtual coronal slice, or if a model projection transverses the virtual slice in a location that is within the voxels assigned to an intermediate structure, between some source and target structure.

**Figure 5:**
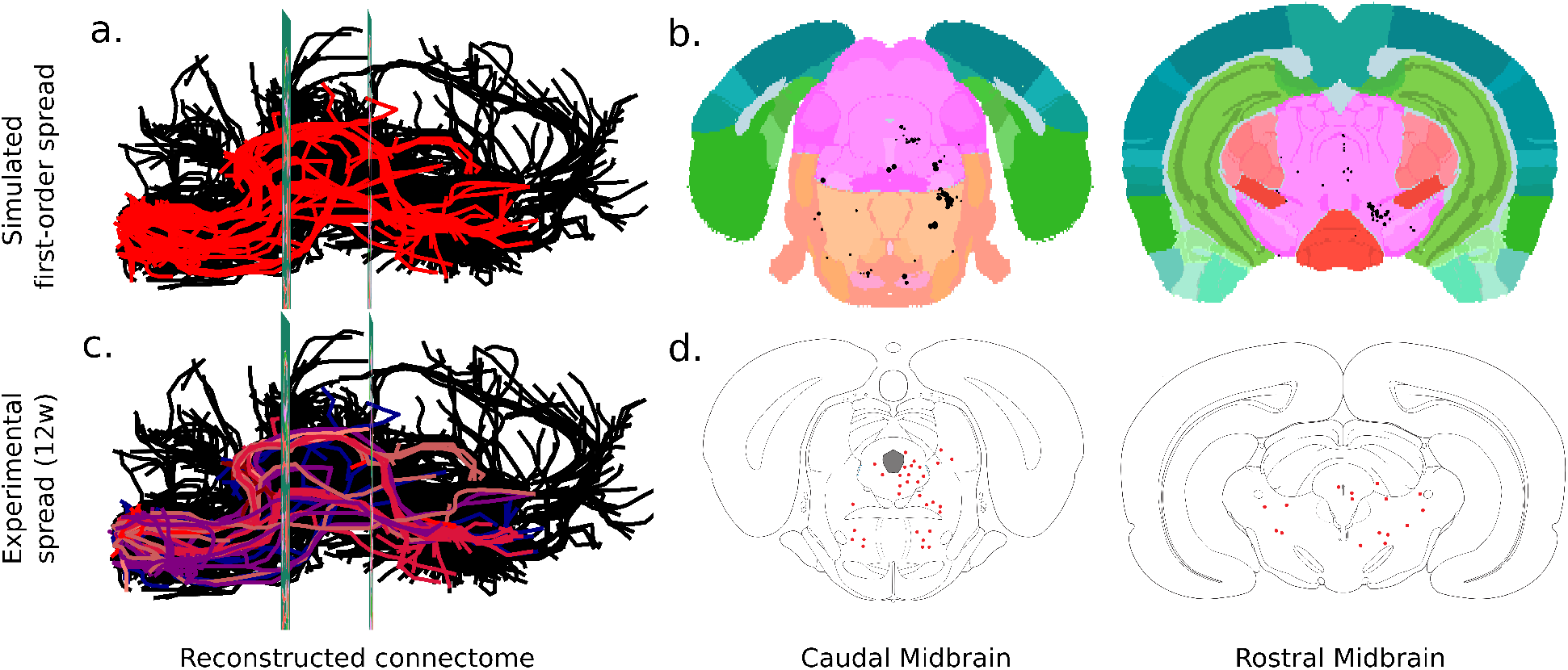
Comparison of simulated and experimental spread. (a) Three dimensional snapshot of the full extent of simulated diffusive spread of virtual *alpha*-synuclein through the first-order mouse connectome (red), started simultaneously from the initiation structures (AMB, DMX). For clarity, only a representative fraction of the non-first-order connectome is illustrated (black). The position of midbrain coronal slices are indicated. (b) Virtual caudal (ABA slice 386, left) and rostral (ABA slice 318, right) midbrain coronal slices intersecting with the first-order mouse connectome (black dots). Increasing rates of simulated spread, in proportion to the number of similar projections (black dots increasing in size). Allen Mouse Brain Atlas structures in standard colours. The virtual coronal brain slices are positioned to correspond to the anatomical position of the microscopically evaluated coronal slices from the *alpha*-synuclein spreading experiments in rats (Ulusoy, Rusconi, et al. 2013). (c) Lateral view of the same parts of the connectome in (a), with the first-order connectome labelled by the severity of spreading in the rat experiments, at 12 weeks post injection (same severity colour coding as in Figure 4). (d) Experimental coronal slices with histologically confirmed sites of *alpha*-synuclein pathology (red dots).

The spread of virtual *alpha*-synuclein along the first-order connectome, within a pair of virtual coronal slices, which are anatomically close to those selected for the histopathological evaluation, is illustrated in Figure 5b. The virtual coronal slices, of both caudal and rostral midbrain, illustrate that virtual *alpha*-synuclein accumulates in first-order neuronal projections that are close to, or reach, brain structures of the medulla, pons, midbrain, hippocampal formation and cerebral cortex. More specifically, the diffusive spread of virtual *alpha*-synuclein reaches the caudal midbrain virtual coronal section (slice 386) within the dorsal raphe, followed by structures of the pons, such as the tegmental reticular nucleus (TRN) and superior central nucleus raphe (CSm), structures of the hippocampal formation, such as the entorhinal area (ENT), and other midbrain structures, such as the cuneiform nucleus and the midbrain trigeminal nucleus. Within the rostral midbrain virtual coronal section (slice 318), virtual *alpha*-synuclein spreads to midbrain structures, such as the SNc and pretectal region (MPT), hypothalamic structures (PH) and olfactory areas (TR3), followed by the CA3 field and entorhinal area of the hippocampal formation, structures of the amygdalar nucleus (BMA), thalamus, hypothalamus (MM) and the red nucleus (RN) of the midbrain.

At 12 weeks post injection, the rat midbrain coronal slices revealed *alpha*-synuclein pathology within many, but not all, of the brain structures predicted to have virtual *alpha*-synuclein accumulation in the *in silico* model. Some brain structures with virtual *alpha*-synuclein accumulation did not display *alpha*-synuclein pathology in the rat model, for example, the SNc, the hippocampal formation and olfactory areas.

### Simulated spread of first-order virtual *alpha*-synuclein

Previously, we assumed that a brain structure, within a virtual coronal slice, was reached by virtual *alpha*-synuclein that structure was a source, target or intermediate brain structure along a model projection. This overestimates the amount of brain structures reached by virtual *alpha*-synuclein in a given coronal slice and ignores brain structures that only intersect with other coronal slices. Therefore, we restricted our assumption and designated a structure as affected if virtual *alpha*-synuclein reached it via a source or target structure, ignoring the intersection of model projections with intermediate brain structures. Under this restricted assumption, we simulated the diffusive spread of virtual *alpha*-synuclein, along the first-order mouse connectome, starting simultaneously from the same two initiation structures, DMX and AMB. Figure 6a compares the first occurrence of virtual *alpha*-synuclein with the severity of *alpha*-synuclein pathology in the rat model.

After initiation within the DMX and AMB, at time-step *t*_7_, virtual *alpha*-synuclein reaches two first-order brain structures, the nucleus of the solitary tract (NTS) and the hypoglossal nucleus (XII). Subsequently, between *t*_8_ and *t*_13_, virtual *alpha*-synuclein reaches multiple other medullary brain structures, only a few of which display *alpha*-synuclein pathology in the rat model, such as, the parvicellular reticular nucleus (PARN), the paragigantocellular reticular nucleus (PGRN) and the caudal part of the spinal nucleus of the trigeminal nerve (SPVC). The parabrachial nucleus (PB), which is severely affected by *alpha*-synuclein pathology in the rat model, is reached by virtual *alpha*-synuclein at *t*_14_, together with other brain structures that have moderate pathology, such as PSV. The remaining medullary and pontine brain structures that display *alpha*-synuclein pathology in the rat model, such as the intermediate reticular nucleus (IRN), pontine central gray (PCG) and the locus ceruleus (LC), display virtual *alpha*-synuclein between *t*_15_ and *t*_19_. Also, other midbrain structures are reached by virtual *alpha*-synuclein within this period, including the midbrain reticular nucleus (MRN) at *t*_17_,which has mild *alpha*-synuclein pathology in the rat model.

**Figure 6:**
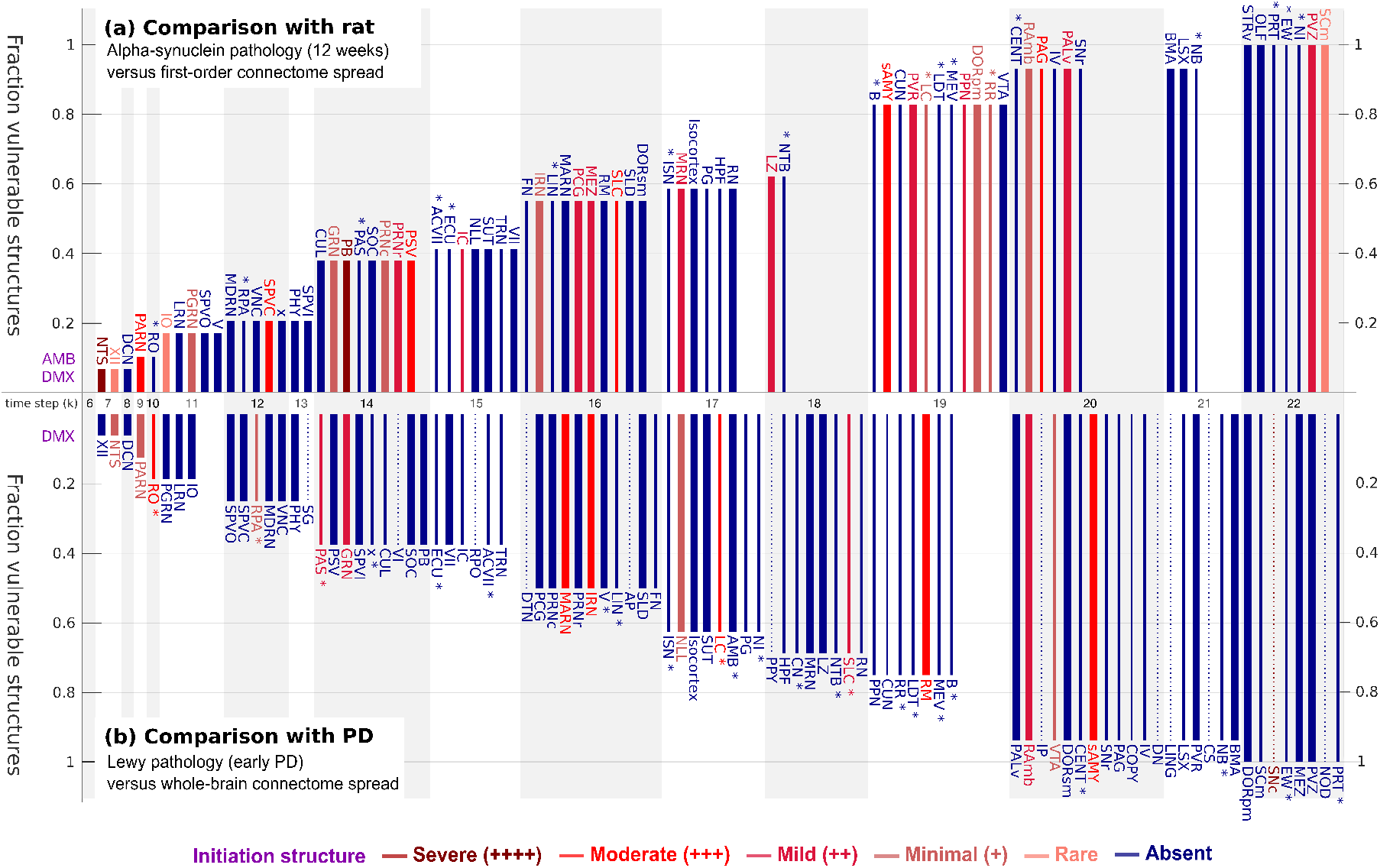
Simulated spread compared with *alpha*-synuclein pathology in rat and early-stage Parkinson’s disease patients. (a) Simulated spread of *alpha*-synuclein, along the first-order connectome, initiated in both the dorsal motor nucleus of the vagus (DMX) and nucleus ambiguus (AMB). The abscissa orders brain structures by the time-step of the first occurrence of virtual *alpha*-synuclein accumulation. The colour of each bar represents the degree of *alpha*-synuclein accumulation observed in the corresponding rat brain structure at 12 weeks post vagal injection. The thickness of each bar represents the number of connections to a first-order brain structure. The ordinate is the fraction of vulnerable rat brain structures, that is, the number of homologous mouse brain structures reached by virtual aggregates at the corresponding time-step, divided by the total number of rat brain structures that demonstrate *alpha*-synuclein pathology at 12 weeks post vagal injection. Each* denotes a *mouse* brain structure that is only anterogradely connected to an initiation structure. There is no clear correlation between the first occurrence of virtual *alpha*-synuclein accumulation (proportional to neuronal projection length) and the severity of *alpha*-synuclein pathology in rats. However more aggregate-positive first-order rat brain structures have a retrograde connection to an initiation structure, than those with only an anterograde connection to an initiation structure (2/18). See Supplementary Table 1 for details. (b) Simulated spread of *alpha*-synuclein spread, along the whole-brain connectome, initiated only in the dorsal motor nucleus of the vagus (DMX). The abscissa orders brain structures by the time-step of the first occurrence of virtual *alpha*-synuclein accumulation. The colour of each bar represents the severity of post-mortem Lewy pathology observed in the corresponding human structure in Parkinson’s disease patients. The ordinate is the fraction of vulnerable human brain structures, that is, the number of mouse brain structures reached by virtual aggregates at the corresponding time-step, divided by the number of homologous human brain structures that have been reported to be vulnerable to degeneration in early PD (Oliveira et al. 2017; Surmeier et al. 2017). Each* denotes a brain structure that is only anterogradely connected to an initiation structure and dashed lines indicate brain structures not directly connected to the DMX. All vulnerable brainstem regions, except the substantia nigra pars compacta (SNc) are directly connected to the dorsal motor nucleus of the vagus (DMX). See Supplementary Table 3 for details.

Supplementary Table 1 provides further detail on the severity of *alpha*-synuclein pathology in the rat spreading model compared with the time-step at which virtual *alpha*-synuclein reaches in the corresponding (target) brain structure within the first-order mouse connectome model. Almost all brain structures affected by *alpha*-synuclein pathology in the rat model are reached by *virtual alpha*-synuclein within three quarters of the simulation steps necessary to cover all first-order brain structures (22 out of 33 time-steps). Most of the brain structures that display *alpha*-synuclein pathology in the rat model are highly connected to the initiation structures (cf Figure 9a). However, there are also many highly connected brain structures that do not display *alpha*-synuclein pathology in the rat model, for example, the superior olivary complex (SOC) and the vestibular nuclei (VNC). Although 18 brain structures receive exclusively anterograde first-order model projections from one of the two initiation structures, only two (LC and RR), display *alpha*-synuclein pathology in the rat model.

**Figure 7:**
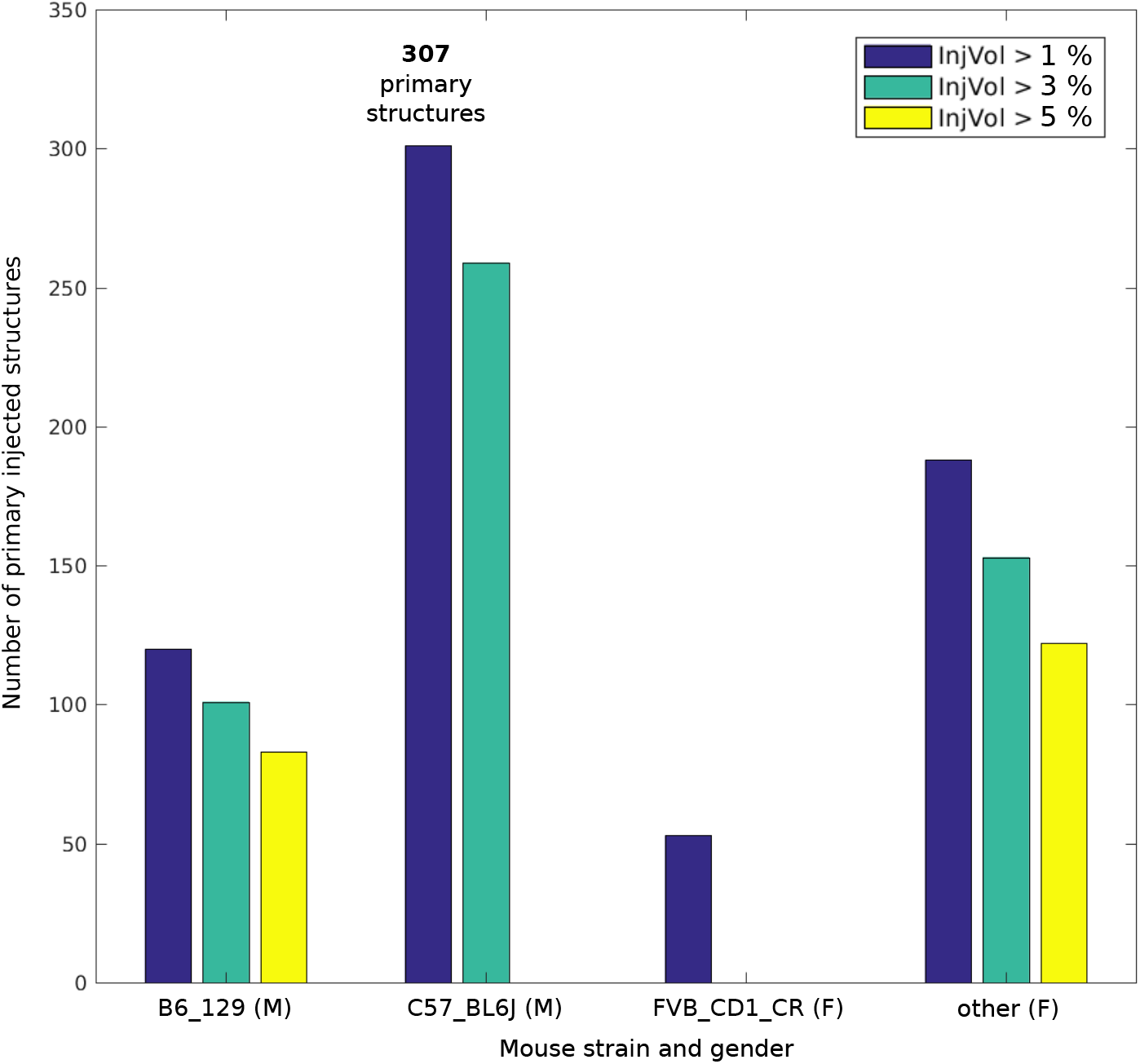
Cerebral anatomical coverage as a function of strain and gender. The number of primary brain structures (ordinate) for each mouse strain and gender (M/F), where at least one experiment included the dorsal motor nucleus of the vagus (DMX) as the primary or a secondary structure, subject to three different cut-offs for the minimum acceptable percentage of voxels in the fluorescently labelled source focus of an experiment that defines a secondary structure. Secondary structures were required to have least 1^Ū^% of the volume of the corresponding primary source structure. The largest number of brain structures injected for the same mouse strain and sex was for the male rat with strain C57BL/6J.

**Figure 8:**
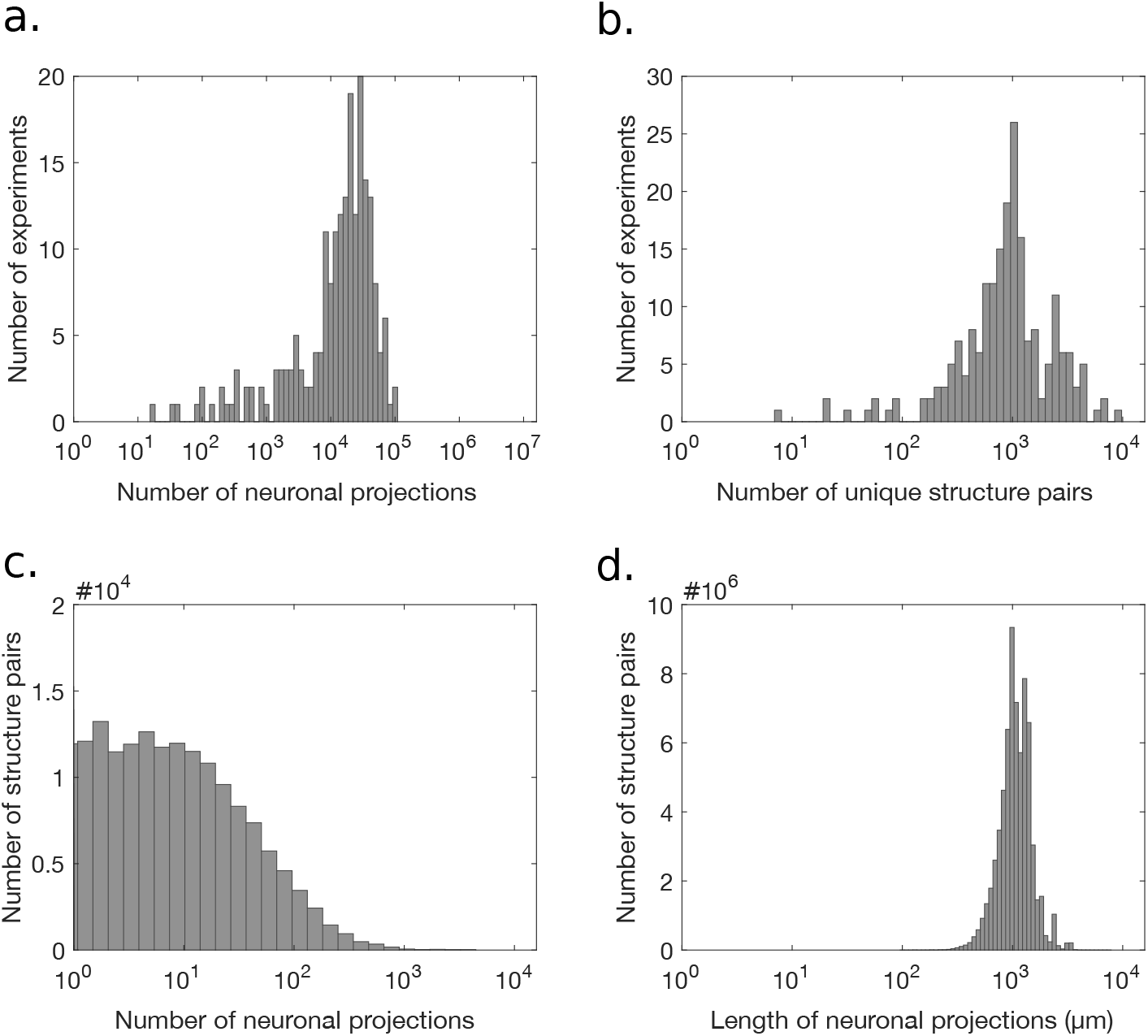
Projection property distributions. The number of neuronal projections (a) and the number of unique structure pairs (b) in each of the 207 selected experiments. Very few experiments have a low number of neuronal projections. The number of neuronal projections (c) and the lengths (*μm*) of neuronal projections (d) in all 207 experiments. Most pairs of connected structures are connected by less than 100 projections. Most neuronal projections are between 500-5000*μm* in length.

**Figure 9:**
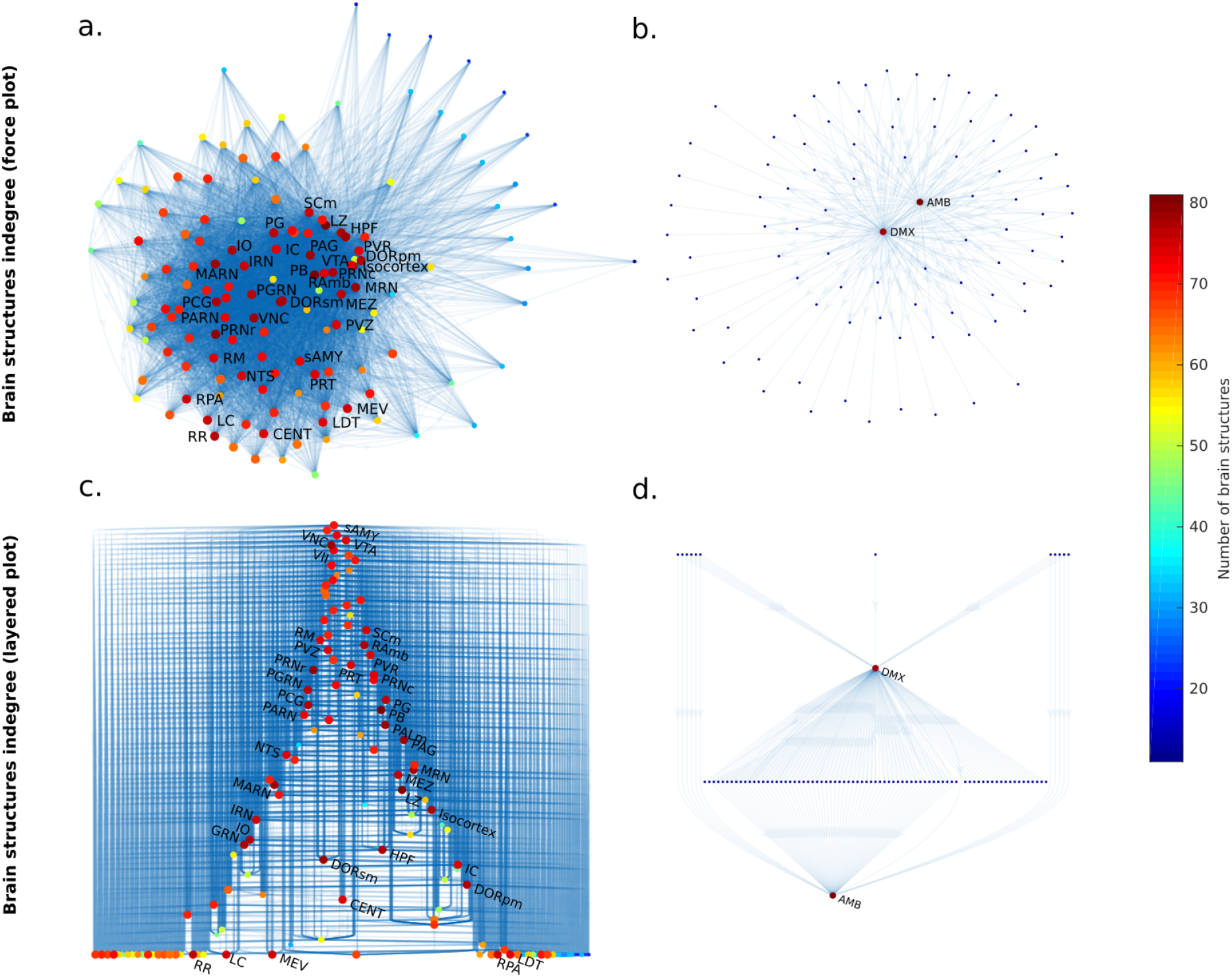
Planar representations of the whole-brain and first-order composite mesoscale mouse connectomes. The first-order connectome considers the dorsal motor nucleus of the vagus (DMX) and nucleus ambiguus (AMB) as initiation structures, all other structures are directly connected to these two initiation structures by one model projection. Each node is a brain structure and each directed edge an anterograde model projection. Each brain structure is colour coded according to the number of source brain structures for which it is a target (indegree). (a, b) Force directed layout of the indegrees of the whole-brain (a) and first-order (b) mesoscale connectome. Attractive forces between highly connected nodes and repulsive forces between poorly connected nodes. Many brainstem structures are highly connected (e.g. NTS, DMX), therefore, selection of the brainstem initiation structure has a large effect on the possible set of first-order structures. (c, d) Directed hierarchical layout of the indegrees of the whole-brain (c) and first-order (d) mesoscale connectome. In the whole-brain, no clear hierarchy is evident. In the first-order connectome, most non-initiation structures are connected to both initiation structures.

### Comparison with Lewy pathology in early Parkinson’s disease

Using the same *in silico* model assumptions, restricted to source and target structures, we also simulated the evolution of diffusive spread of virtual *alpha*-synuclein, initiated only in the dorsal motor nucleus of the vagus nerve (DMX), throughout the whole-brain, composite, mesoscale, mouse connectome. We compared the spread of virtual *alpha*-synuclein with the evolution, distribution and degree of Lewy pathology reported in the brains of patients with early Parkinson’s disease (Surmeier et al. 2017; Oliveira et al. 2017).

As illustrated in Figure 6b, and Supplementary Figure 12, after initiation, virtual *alpha*-synuclein first reaches the hypoglossal nucleus (XII) and the nucleus of the solitary tract (NTS). The NTS has previously been described as having minimal Lewy pathology in Parkinson’s disease and, herein, is followed by other brain structures with minimal and moderate pathology, the parvicellular reticular nucleus (PARN) and nucleus raphe obscurus (RO), respectively. Between *t*_11_ and *t*_19_, multiple medullary and pontine brain regions are reached by virtual *alpha*-synuclein, although only a subset of brain regions have been described as having pathology in early Parkinson’s disease, such as the nucleus raphe pallidus (RPA), intermediate reticular nucleus (IRN), locus ceruleus (LC) and nucleus raphe magnus (RM). Affected midbrain structures in Parkinson’s disease were also reached by virtual *alpha*-synuclein after *t*_18_, with special relevance to the ventral tegmental area (VTA, minimal pathology), midbrain raphe (RAmb, mild pathology) at *t*_20_ and the substantia nigra pars compacta (SNC, severe pathology) at *t*_22_.

According to the mouse connectome, most of the human brain structures with Lewy pathology in Parkinson’s disease are first-order brain structures relative to the DMX, except the substantia nigra pars compacta (SNC) (cf Figure 6b). Furthermore, neither the number of neuronal projections nor the time-step when a structure is first reached by virtual *alpha*-synuclein correlate with the severity of Lewy pathology. The DMX connects to a total of 23 brain structures, only by anterograde first-order model projections, however only 5 (RO, RPA, PAS, LC, SLC) display Lewy pathology *in vivo*. Supplementary Table 3 provides further detail on the severity of Lewy pathology in Parkinson’s disease with the time-step at which virtual *alpha*-synuclein first reaches the corresponding structure within the whole-brain mouse connectome model.

## Discussion

In the rat model we consider, exogenous human *alpha*-synuclein is expressed and transported intracellularly within initiation structures. In contrast to other proteins like GFP, *alpha*-synuclein can be transported from one neuron to another (Ulusoy, Rusconi, et al. 2013; Helwig et al. 2016; Ulusoy, Phillips, et al. 2017; Rusconi et al. 2018). It is due to this specific property of *alpha*-synuclein that we observe exogenous human *alpha*-synuclein in directly connected neurons (first-order connectivity) 12 weeks after rAAV injection (Ulusoy, Rusconi, et al. 2013). It is important to note that this observation of *alpha*-synuclein in recipient neurons is not due to the transport of the virus to the recipient neurons. Using a number of methods, including the use of control virus (rAAV-GFP), RT-PCR analysis of viral genome products in the brain tissue and detection of viral mRNA using a highly sensitive *in situ* hybridization, earlier studies have shown that rAAV virus used at our titer range and injection conditions do not spread trans-cellularly, but it is in fact the protein product (i.e. *alpha*-synuclein) that is transported from one neuron to another (Ulusoy, Rusconi, et al. 2013; Helwig et al. 2016; Ulusoy, Phillips, et al. 2017).

Detection of the exogenous *alpha*-synuclein in rats was performed using a human specific *alpha*-synuclein antibody (i.e. syn211), which does not bind to murine *alpha*-synuclein (Waxman et al. 2008). This enabled us to follow the spread of *alpha*-synuclein unambiguously with high sensitivity. Syn211 binds to the C-terminus of the protein and is capable of detecting monomeric and multimeric forms of *alpha*-synuclein (El-Agnaf et al. 2006). Although the detection method used in this study does not differentiate between *alpha*-synuclein species, using a combination of methods including conformation specific antibodies against oligomeric and fibrillar *alpha*-synuclein, proximity ligase assay and thioflavin S binding, earlier studies showed that the initiation structures contain monomeric, oligomeric and fibrillar forms of *alpha*-synuclein, whereas recipient neurites contain only monomeric and oligomeric forms of *alpha*-synuclein (Helwig et al. 2016).

rAVV injection into the vagus nerve leads to *alpha*-synuclein overexpression in the efferent vagal projections and soma of neurons in the initiation structures, that is, the dorsal motor nucleus of the vagus (DMX) and the nucleus ambiguus (AMB) (Figure 2b). Other brain structures receive afferent vagal projections, e.g., the nucleus of the solitary tract (NTS), and also contain transduced neurites in the rat spreading model. However, when selecting initiation structures for first-order virtual spread, we included structures containing soma within the brain that send efferent projections into the vagus (DMX and AMB), but omitted structures where the soma is outside of the central nervous system and sends afferent vagal projections to the brain (NTS, AP, SPVC). It is important to note that the NTS, AP and SPVC send projections to the DMX and AMB, therefore they are still part of the first-order connectome. This may account for some of the *alpha*-synuclein immunoreactivity observed in within rat NTS, AP and SPVC. Since fibrillar forms are found in initiation structures but not other structures, it would be interesting to compare the molecular structure and post-translational modifications of neuritic *alpha*-synuclein as a result of viral transduction, versus somal *alpha*-synuclein as a result of first-order retrograde spread, within any one of the latter three structures in future studies.

Is connectivity necessary to determine which brain structures will display *alpha*-synuclein pathology? Starting from the initiation structures (DMX, AMB), our first-order mesoscale mouse connectome includes all except one of the microscopically resolved rat brain structures with *alpha*-synuclein pathology *in vivo* (Supplementary Table 1). The one exceptional structure was the area postrema (AP, cf Figure 2), which is known to receive projections from the inferior ganglion of the vagus (IGX) (Gwyn et al. 1979) and to send projections to the initiation structures (DMX, AMB) (Shapiro et al. 2004). However, to date, the area postrema has never been a source structure in a published Allen Mouse Connectivity experiment, and neither of the initiation structures send projections to the area postrema (AP), therefore its exclusion from a first-order connectome, derived only from Allen Mouse Connectivity Data, is expected. Therefore, we conclude that the connectome is necessary for spread of induced *alpha*-synuclein pathology in rats.

Is connectivity sufficient to determine which brain structures will display *alpha*-synuclein pathology? Our quantitative simulations predict highly anatomically resolved patterns of both retrograde and anterograde spread of virtual *alpha*-synuclein through the connectome and thereby predict the brain structures that *could* display *alpha*-synuclein pathology induced from connected neuronal structures, but not the structures that *will* display *alpha*-synuclein pathology (Figures 5 and 6a). Many first order brain structures displayed accumulation of virtual *alpha*-synuclein but do not display any *alpha*-synuclein immunoreactivity in the rat spreading model, e.g., SOC, VNC, medullary reticular nucleus (MDRN), nucleus raphe obscurus (RO), intermediate reticular nucleus (IRN), magnocellular reticular nucleus (MARN) and nucleus raphe pallidus (RPA). Therefore, we conclude that the connectome is not sufficient for spread of induced *alpha*-synuclein pathology in rats.

Does the amount, length, or direction, of neuronal projections determine the brain structures that will display *alpha*-synuclein pathology? Most of the brain structures that display *alpha*-synuclein pathology in the rat spreading model are highly connected to the initiation structures (Figure 6a). This suggest that, for vulnerable neurons, the propensity for spread between two nuclei increases when there are more neuronal projections between them. Almost all brain structures with *alpha*-synuclein spreading *in vivo* are reached by virtual *alpha*-synuclein before three quarters of the simulation time-steps necessary to cover all first-order brain structures. This suggests that, for vulnerable neurons, spread may be more efficient for shorter projections lengths. However, there are also many examples of structures that are highly connected to the initiation structures by relatively short projections, but do not show *alpha*-synuclein pathology in the rat spreading model. Amongst the 29 first-order connectome brain structures that show *alpha*-synuclein pathology *in vivo*, 27 send projections to an initiation structure, while only 2 receive projections from an initiation structure (LC and RR). This suggest that the spread of pathology is much more likely to occur in a retrograde than anterograde direction. To refine these conclusions, it would be ideal to examine cleared brains (Richardson et al. 2015) from a rat cohort sacrificed at a regular intervals post viral transduction, with a subset of the cohort that have had specific neuronal connections ablated a priori.

In comparison with direct injection into brain structures, a vagal injection allows a more controlled initiation of exogenous *alpha*-synuclein expression within a set of known brain structures. This is a key consideration when studying the hypothesis that Lewy pathology spreads via the dorsal motor nucleus of the vagus. Nevertheless, this injection method still allows a certain degree of uncertainty in the distribution of viral expression of *alpha*-synuclein, albeit in a limited number of brainstem structures (DMX, AMB). It would be beneficial to develop rodent models where it is certain that only one brainstem structure could express and spread *alpha*-synuclein to differentiate between retrograde and anterograde transmission, for example by using tissue specific promoters or conditional expression models (i.e. cre-lox systems). This would permit more precise initiation of the computational model. Furthermore, histopathological evaluation of *alpha*-synuclein spreading at multiple time points, and at mesoscale resolution, would permit a more refined understanding of the relative resistance versus vulnerability of brain structures, within the context of the neuronal connectome. Nevertheless, we believe that we have already sufficient resolution to make firm conclusions about the patterns of spreading of *alpha*-synuclein pathology in rats.

We compared human pathology with computational modelling of spread along the mouse connectome, even though a comprehensive human cerebral connectome, at single neuronal resolution, would be superior for direct comparison with pathology progression in Parkinson’s disease. However, neuronal anatomy of the brainstem in evolutionarily well conserved in mammals. Assuming that, for the brainstem, the mouse mesoscale connectome is a fair representation of the human connectome, our *in silico* model predicts the contribution of neuronal network topology and geometry alone to the selective vulnerability of brain structures in Parkinson’s disease. The Braak hypothesis posits spread via the DMX, therefore, in our simulation of diffusive spread through the whole-brain connectome, we also initiate spread from the DMX. Interestingly, brain structures known to be vulnerable in early Parkinson’s disease, inclusive of less-vulnerable structures, were predicted to receive spread before the substantia nigra pars compacta (Figure 6 and Supplementary Table 3). In concordance with the observations of Braak et al. (**braak_neuropathological_**), we find that most vulnerable brain-stem structures are indeed directly connected by neuronal projections to the DMX. This observation further supports the interpretation that neuronal connectivity is necessary for spread of *alpha*-synuclein.

Multiple highly vulnerable brain structures in early Parkinson’s disease, such as raphe obscurus (RO) and locus coeruleus (LC), were simulated to receive spread of virtual *alpha*-synuclein earlier than less-vulnerable counterparts, respectively, the raphe pallidus (RPA), sub-coeruleus (SLC), except for the ventral tegmental area (VTA) and substantia nigra pars compacta (SNC). Interestingly, all vulnerable structures associated with prodromal Parkinson’s disease (Surmeier et al. 2017) are reached by virtual *alpha*-synuclein earlier than the substantia nigra pars compacta. However, multiple brain structures known to be resistant to Lewy pathology in Parkinson’s disease, e.g., the hypoglossal nucleus (XII), accumulate virtual *alpha*-synuclein at early simulation time-steps. These observations support the interpretation that the connectome not sufficient to determine selective vulnerability nor to predict the degree of vulnerability.

Apart from neuronal connectivity, multiple other factors, such as alterations in protein degradation, mitochondrial biogenesis and endo- or exocytosis, might contribute to the selective vulnerability of similarly connected brain structures in Parkinson’s disease (Surmeier et al. 2017). Deep phenotyping of connected brain structures with different degrees of vulnerability would increase our understanding of the contribution of these other factors to selective vulnerability. Our *in silico* model simulates the spread of arbitrary aggregates through a static neuronal network, which does not yet dynamically evolve to account for the dynamic deterioration of the neuronal projections that occurs after sufficient *alpha*-synuclein pathology *in vivo*. Although the Allen Brain Institute connectome data can provide resolution of individual neuronal projections at the site of injection, a variably sized focus of fluorescence is observed in raw data and involves the injected structure and a set of the neighbouring structures. Such artefacts may overestimate the size of the source focus sending projections to other brain areas. Therefore, additional connectivity data with more highly resolved structure-specific projection data for certain key brain structures, e.g., DMX or LC (Henrich et al. 2018), might allow a more refined prediction of the contribution of the connectome to the spread of pathology. Nevertheless, we do not believe such data would affect our overall conclusion that the connectome is necessary but not sufficient to explain the patterns of spreading of *alpha*-synuclein pathology in rats.

Many other neurodegenerative diseases target large-scale human brain networks (Seeley et al. 2009; Brettschneider et al. 2015) and, similarly to Parkinson’s disease, multiple mouse and rat models are being developed to study pathology progression and protein spreading (Brettschneider et al. 2015). Our whole-brain connectome model was first generated to comprehensively represent anatomy, and thereafter, selectively applied to model Parkinson’s disease progression. Therefore, our model could readily be applied to study the contribution of the connectome in the selective progression of pathology in other neurodegenerative diseases, e.g., Alzheimer’s disease, or Machado-Joseph’s disease.

## Conclusions

We simulated the contribution of the cerebral connectome to the spread of virtual *alpha*-synuclein using a novel computational model of spatiotemporal spread within a composite representation of the mouse mesoscale connectome. We conclude that neuronal connectivity is necessary but not sufficient to computationally predict the anatomical pattern of induced *alpha*-synuclein pathology observed in rats. We also observe that the spread of *alpha*-synuclein pathology in rats is much more likely to occur in a retrograde than anterograde direction. The propensity for spread between two brainstem nuclei seems to increase the shorter the length of the projection and more neuronal projections there are between them. However, there are also many examples of structures that are highly connected to the initiation structures by relatively short projections that do not display *alpha*-synuclein pathology in a rat spreading model. If one assumes that mouse brainstem connectivity is a fair representation of human brainstem connectivity, then our results suggest that the connectome is necessary, but not alone sufficient, to predict the histopathological patterns of Lewy pathology observed in Parkinson’s disease.

## Nomenclature

AAA: Anterior amygdalar area
Allen SDK: Allen Brain Institute Software Development Kit
AMB: Nucleus ambiguus
AP: Area postrema
APN: Anterior pretectal nucleus
CEAc: Central amygdalar nucleus capsular
DMH: Dorsomedial nucleus of the hypothalamus
DMX: Dorsal motor nucleus of the vagus nerve
GRN: Gigantocellular reticular nucleus
IGX: Inferior ganglion of the vagus (nodose ganglion)
IRN: Intermediate reticular nucleus
LC: Locus ceruleus
LHA: Lateral hypothalamic area
MARN: Magnocellular reticular nucleus
MDRN: Medullary reticular nucleus
MEV: Midbrain trigeminal nucleus
MG: Medial geniculate complex
MRN: Midbrain reticular nucleus
NTS: Nucleus of the solitary tract
PAG: Periaqueductal gray
PARN: Parvicellular reticular nucleus
PB: Parabrachial nucleus
PCG: Pontine central gray
PD: Parkinson’s disease
PG: Pontine gray
PP: Peripeduncular nucleus
PPN: Pedunculopontine nucleus
PRN: Pontine reticular nucleus
PSTN: Parasubthalamic nucleus
PSV: Principal sensory nucleus of the trigeminal
PVH: Paraventricular hypothalamic nucleus
RM: Nucleus raphe magnus
RN: Red nucleus
RO: Nucleus raphe obscurus
RPA: Nucleus raphe pallidus
RR: Midbrain reticular nucleus retrorubral area
SBPV: Subparaventricular zone
SC: Superior colliculus motor related
SGX: Superior ganglion of the vagus (jugular ganglion)
SLC: Subceruleus nucleus
SNc: Substantia nigra pars compacta
STN: Subthalamic nucleus
TRN: Dorsal tegmental nucleus
TU: Tuberal nucleus
XII: Hypoglossal nucleus

## Author contributions

R.F. conceived the project. M.O. and S.A. developed the computational model and analysed the simulation results. G.P. downloaded the mouse connectivity data. A.U. and D.D. obtained and analysed the rat *in vivo* data. M.O., S.A., A.U., D.D. and R.F. wrote the manuscript.

## 1 Supplementary Methods

### 1.1 Allen Brain Atlas connectivity data

In the Allen Mouse Brain Reference Atlas, a *structure* is a neuroanatomical region of interest. Structures are hierarchically organised into an ontology. The corresponding StructureGraph starts with a Root structure, representing the whole-brain. Every other structure is a daughter via a PartOf relationship. Descending the hierarchy, one traverses from lower to higher anatomical resolution, terminating with a *leaf structure*, which is a daughter that is not a parent, and therefore the highest level of structural resolution in the atlas. Each structure is also assigned a consistent colour, which we also adhere to in our visualisation results.

The Allen Mouse Brain Connectivity is a mesoscale mouse connectome (Oh et al. 2014), generated by integration of the data resulting from many separate connectivity experiments. In each connectivity experiment a single stereotaxic injection of an anterograde tracer, recombinant adeno-associated virus expressing a fluorescent protein under the control of a neuron specific Synapsin 1 promotor (rAAV-GFP), is aimed into a single cerebral structure. Serial two-photon tomography is used to acquire fluorescent images and a sophisticated image processing and analysis pipeline is subsequently used to produce anatomically specified neuronal projections, consisting of fluorescent voxels, registered to common coordinates provided by the Allen Mouse Brain Reference Atlas.

Often the neuronal projections from a single connectivity experiment emanate from more than one *source structure*. Even though the injection is aimed toward a single structure, a number of cell bodies in the neighbouring structures may be transduced by rAAV-GFP and emanate fluorescent neuronal projections, due to the diffusion of the viral vector around the point of injection (Figure 2a). As a result, a connectivity experiment may also contain projections that originate from more than one source structure.

Each connectivity experiment consists of information about the strain and gender of the mouse (Figure 1a), about all source structures and about each neuronal projection (Figure 1c). Each efferent neuronal projection is specified by a single source structure, a set of voxel coordinates along the projection, and a single *target structure*, where the projection ends. Each connectivity experiment is identified by a seven-digit number, with prefix `ABAID`, available as lightweight data-interchangeable file `JSON` file (`JavaScript Object Notation`). Our capture and preprocessing of connectivity data was implemented in Python, leveraging the `Allen Software Development Kit` (`Allen SDK`) available from http://alleninstitute.github.io/AllenSDK/ and the Mouse Connectivity Application Programming Interface, available from http://help.brain-map.org/display/mouseconnectivity/API.

### 1.2 Selection of connectivity data

Each connectivity experiment fluorescently labels a three dimensional *source focus* in the brain, the volume of which reflects the amount of fluorescence above a threshold (Oh et al. 2014), as well as a variable amount of neuronal projections emanating from this focus. A focus may overlap with multiple brain structures, that are identified with the highest level of anatomical resolution feasible for that experiment. The *primary* source structure is defined as the structure with the majority of voxels in the source focus. To account for AAV transduction outside the primary source, for each experiment, we also include just two *secondary* source structures with the second and third greatest percentage of voxels in the source focus (Figure 1b and 1d). Importantly, all focus structures were required to belong to different *anatomical branches* and to not have *root* as parent and secondary structures were required to have least 1% of the volume of the corresponding primary source structure (Figure 1 b). The primary and the two secondary source structures, as defined in the Allen Mouse Brain Connectivity data, are used to reconstruct the mouse connectome (Figure 1f and 1j).

The aforementioned selection of experiments did not include connectivity experiments that were excluded after quality control of the projections contained within it (Figure 1h). For example, if the distance between the first two coordinates was more than *3000μm*, that projection was considered spurious and excluded from further analyses. Each connectivity experiment can be summarised as a set of directed links between unique pairs of source and target structures. A total of 10 experiments were excluded from further analyses because the removal of spurious projections reduced the number of quality controlled projections to zero.

The Allen Mouse Brain Connectivity is an amalgamation of more than 2,900 separate connectivity experiments. However, we selected a subset of 207 experiments to reconstruct a composite mesoscale mouse connectome to use for simulation of spread within the whole-brain. This selection corresponded to the largest number of cerebral source structures (439 focus structures) for the same mouse strain (C57BL/6J), from the same sex (male) and containing source structures that were initiation structures in the rat spreading model (Supplementary Figure 7).

To assess the quality ofthe selected experiments, several topological and geometrical properties of the connectivity data from the 207 selected experiments were assessed. Specifically, the number of neuronal projections and number of unique structure pairs in each experiment, as well as the number of neuronal projections and the lengths of neuronal projections.

## 2 Rat model of caudo-rostral *alpha-synuclein* spreading

### 2.1 *Viral* Vectors

Recombinant adeno-associated virus (serotype 2 genome and serotype 6 capsid, rAAV) was used for transgene expression of human wild-type *alpha*-synuclein (Sirion Biotech, Martinsried, Germany). Human alpha-synuclein sequence was placed downstream to human Synapsin 1 promoter. Gene expression was enhanced by placing a woodchuck hepatitis virus post-transcriptional regulatory element and a polyadenylation signal sequence downstream to the gene sequence (Ulusoy, Rusconi, et al. 2013).

### 2.2 Animals and surgical procedure

Four young adult female Sprague Dawley rats weighing 200–250 g were obtained from Charles River (Kisslegg, Germany). They were housed under a 12-h light/12-h dark cycle with free access to food and water. Experimental design and procedures were approved by the ethical committee of the State Agency for Nature, Environment and Consumer Protection in North Rhine Westphalia. 2 μL of (5 × 10^12^ genome copies/mL) vector solution was injected into the left vagus nerve at the level of the neck, distal to the nodose ganglion, as described earlier (Ulusoy, Musgrove, et al. 2015; Rusconi et al. 2018).

### 2.3 Tissue preparation

Animals were killed under pentobarbital anaesthesia 12 weeks after the virus injection and perfused through the ascending aorta with saline, followed by ice-cold 4% (w/v) paraformaldehyde. Brains were immersion-fixed in 4% paraformaldehyde (for 24 h) and transferred into 25% (w/v) sucrose solution for cryopreservation. 40 μm sections were cut on a freezing microtome in the coronal plane.

### 2.4 Histology

For histological analysis, free-floating coronal sections were stained against human alpha-synuclein as described earlier (Ulusoy, Rusconi, et al. 2013). Briefly, brains were treated with mouse anti-human-*alpha*-synuclein clone syn211 (36– 008, Merck Millipore; 1:10,000 dilution) overnight at room temperature followed by 1 hour incubation in biotinylated secondary antibody solution (horse anti-mouse BA2001; Vector Laboratories, Burlingame, CA, USA). Following treatment with avidin-biotin-peroxidase (ABC Elite kit, Vector Laboratories), sections were developed using 3,3’-diaminobenzidine kit (Vector Laboratories).

Examination of stained sections were carried out with an Axioscope microscope using a 40x Plan-Apo oil objective (Zeiss). The 829 leaf structures listed in the Allen Mouse Brain Atlas were simplified into 139 rat brain structures, that were resolved by epiflourescent microscopy of brain sections, by merging subnuclei that were not unambiguously separable into single nuclei. For example, the seven subnuclei of the parabrachial nucleus (i.e. lateral division dorsal lateral part, lateral division central lateral part, lateral division external lateral part, lateral division ventral lateral part, medial division medial medial part, medial division external medial part and lateral division superior lateral part) were merged and analysed as a single structure, namely parabrachial nucleus. Similarly, for example, the central amygdalar nucleus was analysed as a single structure rather than medial, lateral and capsular subnuclei. However, any subnuclei that were unambiguously visually separable were kept as separate structures, for example the substantia nigra pars compacta and pars reticulata.

The presence of human *alpha*-synuclein containing neurites was investigated in every-sixth section throughout the entire brain in the 139 aforementioned microscopically resolved rat brain structures (Paxinos et al. 2007). Human *alpha*-synuclein containing neurites appeared as antibody labelled fine threads, often containing enlarged bead-like varicosities (Ulusoy, Rusconi, et al. 2013). The number of human *alpha*-synuclein immunoreactive neurites were counted for each brain structure at the section corresponding to the Bregma coordinate where the structure is at its largest size (Paxinos et al. 2007). The average number of human *alpha*-synuclein-immunoreactive neurites were used to define a severity score according to the following parameters: - (absent): 0 neurites; rare: <2 neurites; + (minimal): 2-5 neurites;++ (mild): 6-10 neurites; +++ (moderate): 10-20 neurites and ++++ (severe): >20 neurites. This semi-quantitative data (Figure 1l) were compared with quantitatively predictions of virtual *alpha*-synuclein accumulation at each brain structure independently (Figure 1m).

### 2.5 Computer code

Custom computer code accompanies this manuscript allowing one to completely reproduce every step of the pipeline, from data collection to visualisation, including scripts for accessing the Allen API, preprocessing the data and visualisation. (Comment to reviewers and editors: Upon request, this code will be made available to editors and referees, via a version controlled repository.)

## 3 Supplementary Results

### 3.1 Analysis of Allen Brain Atlas connectivity experiments

The Allen Mouse Brain Connectivity Atlas is an amalgamation of more than 2,900 separate connectivity experiments. However, we selected a subset of 207 of these experiments to reconstruct a composite mesoscale mouse connectome to use for simulation of spread within the whole-brain. This selection corresponded to the largest number of cerebral source structures (307 primary structures out of 439 focus structures) for the same mouse strain (C57BL/6J), from the same sex (male) and containing source structures that were initiation structures (DMX, AMB) of the spread of alpha-synuclein pathology in the rat model (Supplementary Figure 7).

The selected set of 207 connectivity experiments contains 307 primary injected structures and 132 secondary injected structures which correspond to almost all (829/836) leaf structures in the whole mouse brain. After quality control for spurious projections, an average connectivity experiment contained 21, 338 individual neuronal projections, but in average it represents connections between only 1, 277 unique pairs of brain structures (Supplementary Figure 8c). Most of the experiments have around 10^4^ – 10^5^ neuronal projections that correspond to 10^3^ unique structure pairs (Supplementary Figure 8a-b). On average, a neuronal projection within the selected experiments (after quality control for spurious projections) have 10^3^ *μm* of length (Supplementary Figure 8d). This analysis also ensured that no experiment, with primarily short neuronal projections, was included. Such experiments were eliminated from consideration as they were interpreted as non-specific fluorescent labelling.

### 3.2 Topological analysis of the whole-brain and first-order connectomes

Supplementary Figure 9 provides a visual perspective on the topological properties of the whole-brain and first-order connectomes, while Supplementary Figure 10 places these topological properties in their anatomical context.

The first-order connectome is a small subset of the whole-brain connectome that contains only brain structures that are directly connected to the initiation structures (DMX and AMB). The direction of each model projection is set by convention to follow an anterograde neuronal projection direction, from a somal source to a neurite terminating in a target structure. The *indegree* of a structure is the number of other structures that send a model projection to it. The *outdegree* of a structure is the number of other structures that it sends a model projection to. Neither the indegree or outdegree are not weighted by the number of similar neuronal projections that correspond to one model projection. Within the brain, there is a widely varying degree of connectivity for any given structure, but most brainstem structures are connected to a large number of other structures. Furthermore, even if a brain structure in the whole-brain connectome, or first-order connectome, is highly connected, there can be a large imbalance between its indegree and outdegree. For example, the DMX mainly projects neurons into the vagus, but the vagal terminations are outside the brain, so its outdegree in the whole-brain connectome is low. However, the DMX receives a lot of projections from other parts of the brain, so its indegree is high.

**Figure 10:**
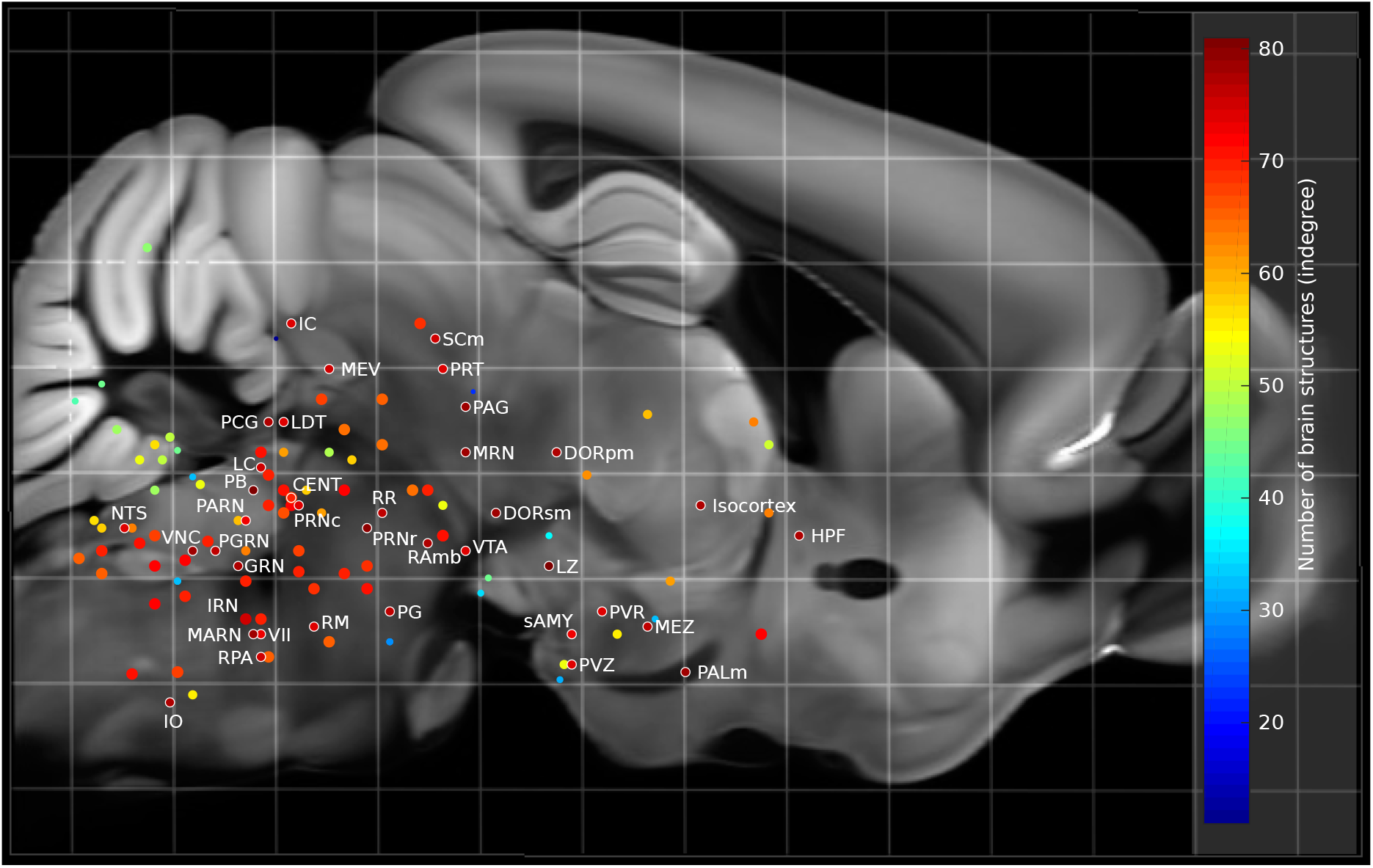
Indegree of microscopically resolved rodent structures. Lateral projection of mouse brain structures (disks) onto a mid-sagittal cerebral section. Disks are colour coded by indegree in the whole-brain connectome, however only those mouse structures that correspond to a microscopically resolved structure in the rat brain spreading experiments are displayed. The high indegree of most rodent brainstem structures is evident.

### 3.3 Evolution of simulated spread

**Figure 11:**
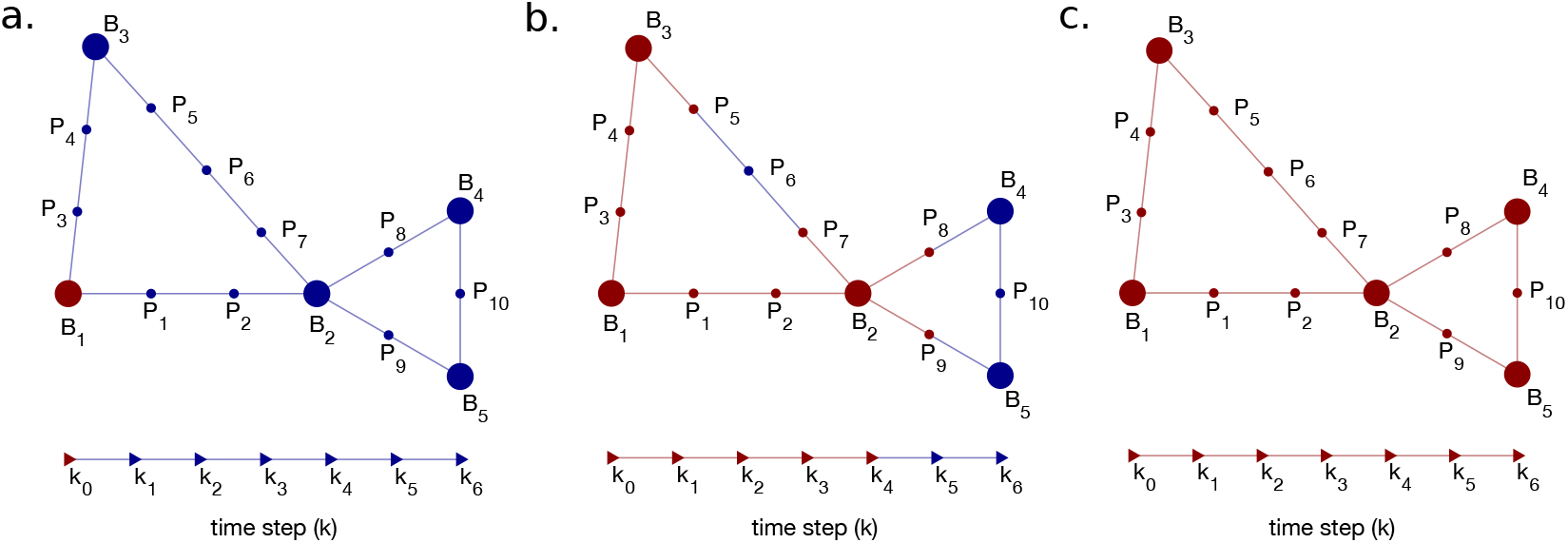
Evolution of diffusive simulated spread. Discretisation of projections and naming conventions for the state variables in a toy network. Path vertices are labelled *P_i_* and brain structure vertices are labelled as *B_i_*. (a) At time step *k*_0_, brain structure *B*_1_ is initiating the diffusive spread (red) on a small network (blue). (b. and c.) Evolution of the diffusive spread (red) between time steps *k*_0_ and *k*_6_. In (b), note that *P*_5_ and *P*_7_ become positive for exogenous alpha-synyclein at time-step *k* = 4 since both are a total of 4 edges in distance away from *B*_1_.

**Figure 12:**
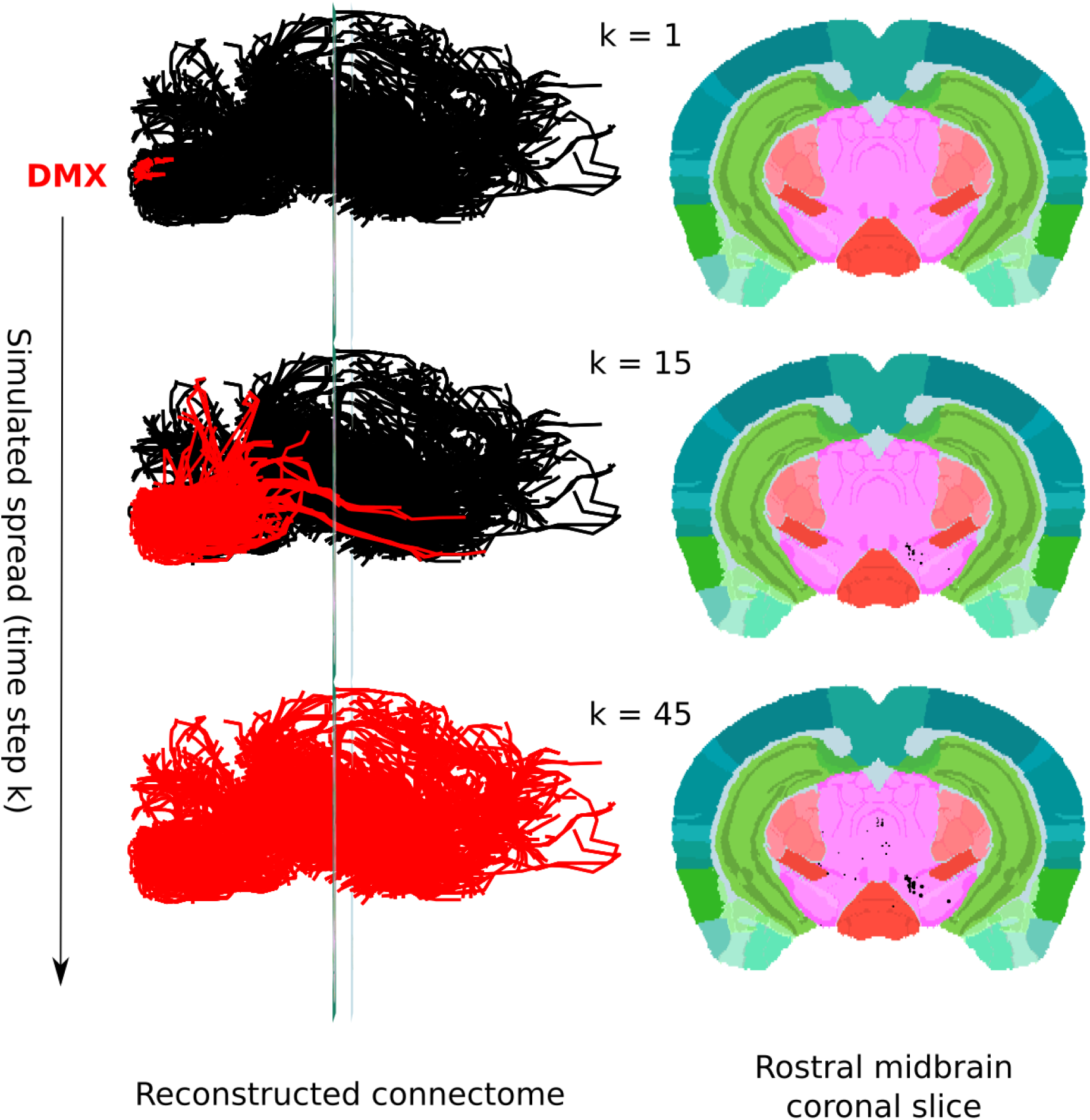
Evolution of simulated spread within the whole-brain mouse connectome. In this simulation, the spread of virtual *alpha*-synclein was initiated within the dorsal motor nucleus of the vagus. In the 3D view (left), the connectome is illustrated via a set of paths (black) each of which is one model projection. Virtual *alpha*-synclein accumulation in both path vertices, at either end of a path edge, is illustrated by highlighting the corresponding path edge in red. Hence, the spread of virtual*alpha*-syndein with respect to time is represented with a dynamically spreading red colour along the path, from dorsal motor nucleus of the vagus throughout the connectome along a sequence of time-steps (k). In the 2D mouse coronal brain slice view (right), each black disc represents virtual *alpha*-syndein accumulation within a model projection path, or within a specific leaf brain structure, that intersects the coronal slice at that position. The diameter of the disc is proportional to the amount of virtual *alpha*-synclein accumulation at that time step. Colours in the simulated midbrain slice (ABA slice 300) are in accordance with the Allen Mouse Brain Reference Atlas. Note that, to simplify the visualisation of spread in 3D, only the projections in the first-order connectome plus the additional projections from connectivity experiments that sent a projection to either the DMX or AMB are shown. However, the corresponding 2D coronal slices show the spread of virtual *alpha*-synclein along the whole-brain mouse connectome.

### 3.4 Detailed comparison between first-order mouse simulation and *alpha*-synuclein spreading in rats

**Table 1:**
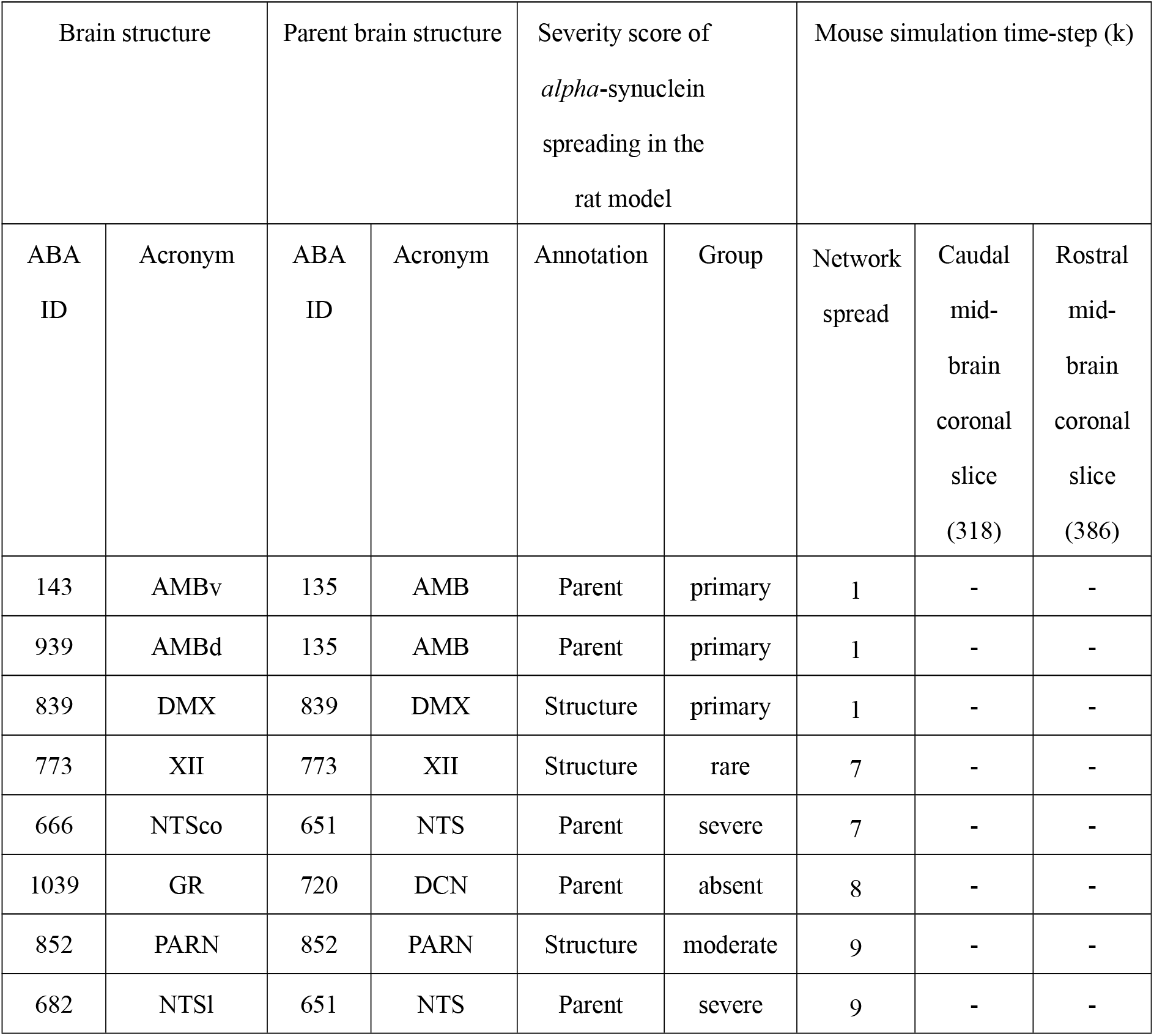
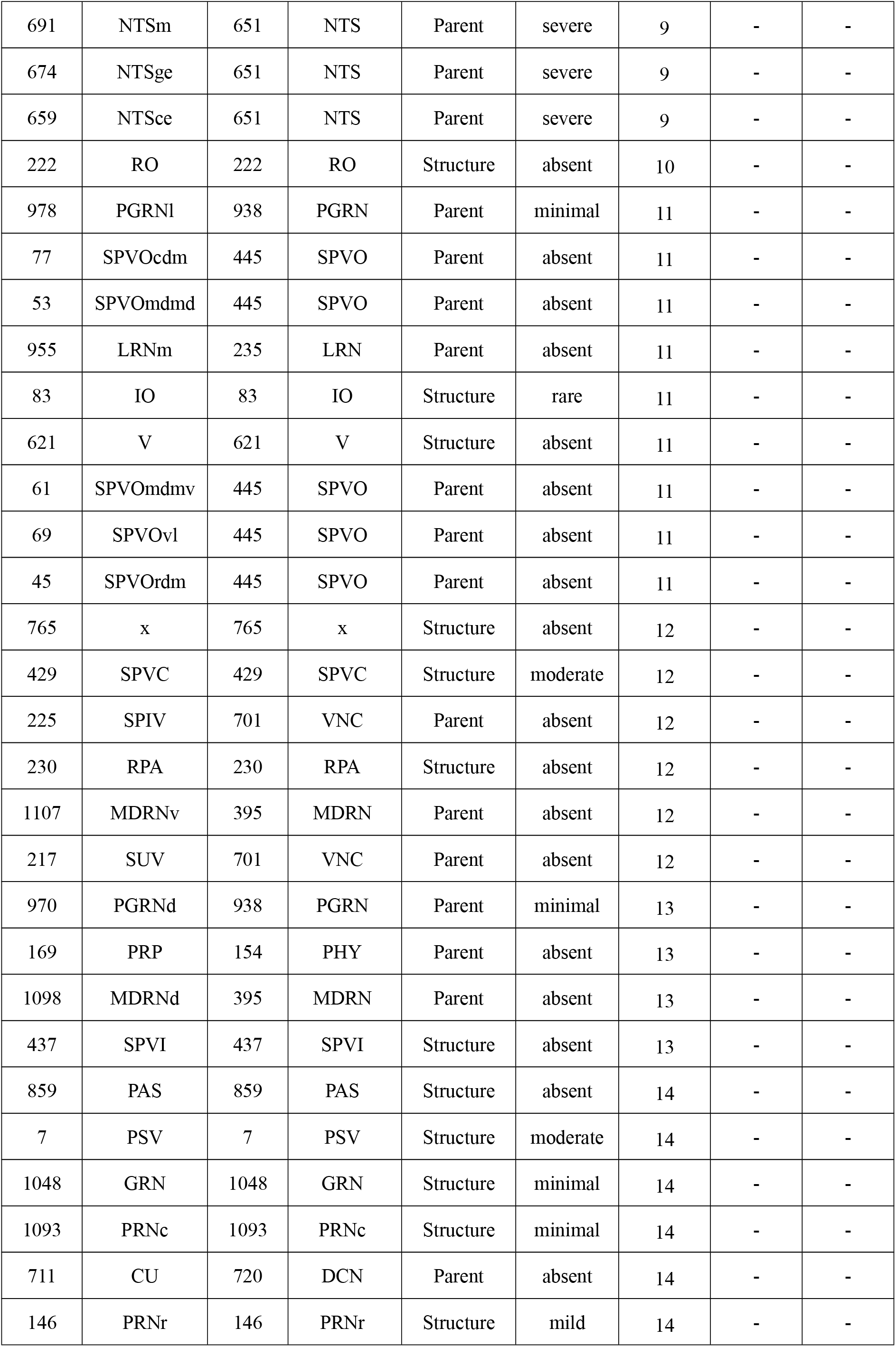
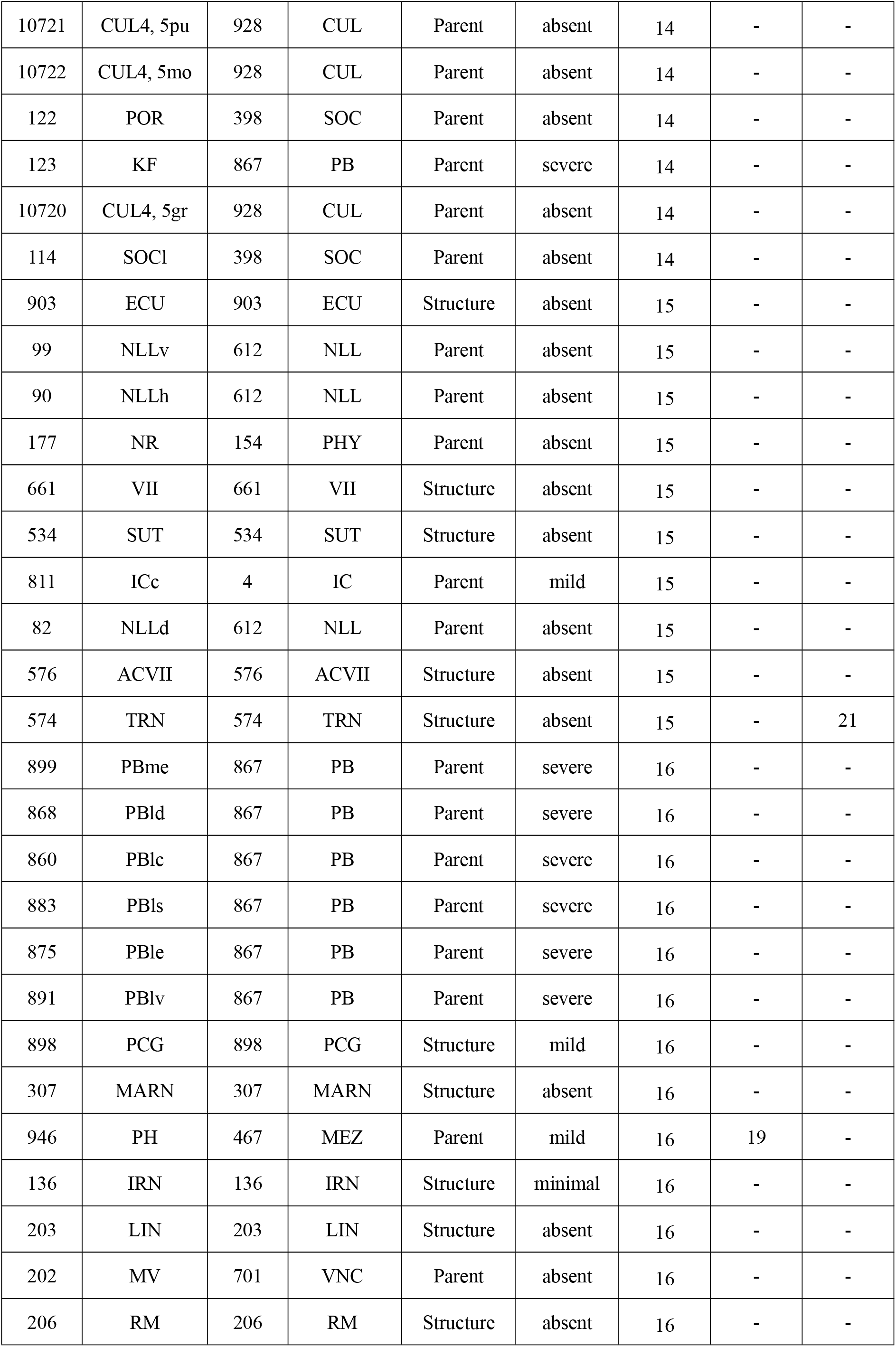
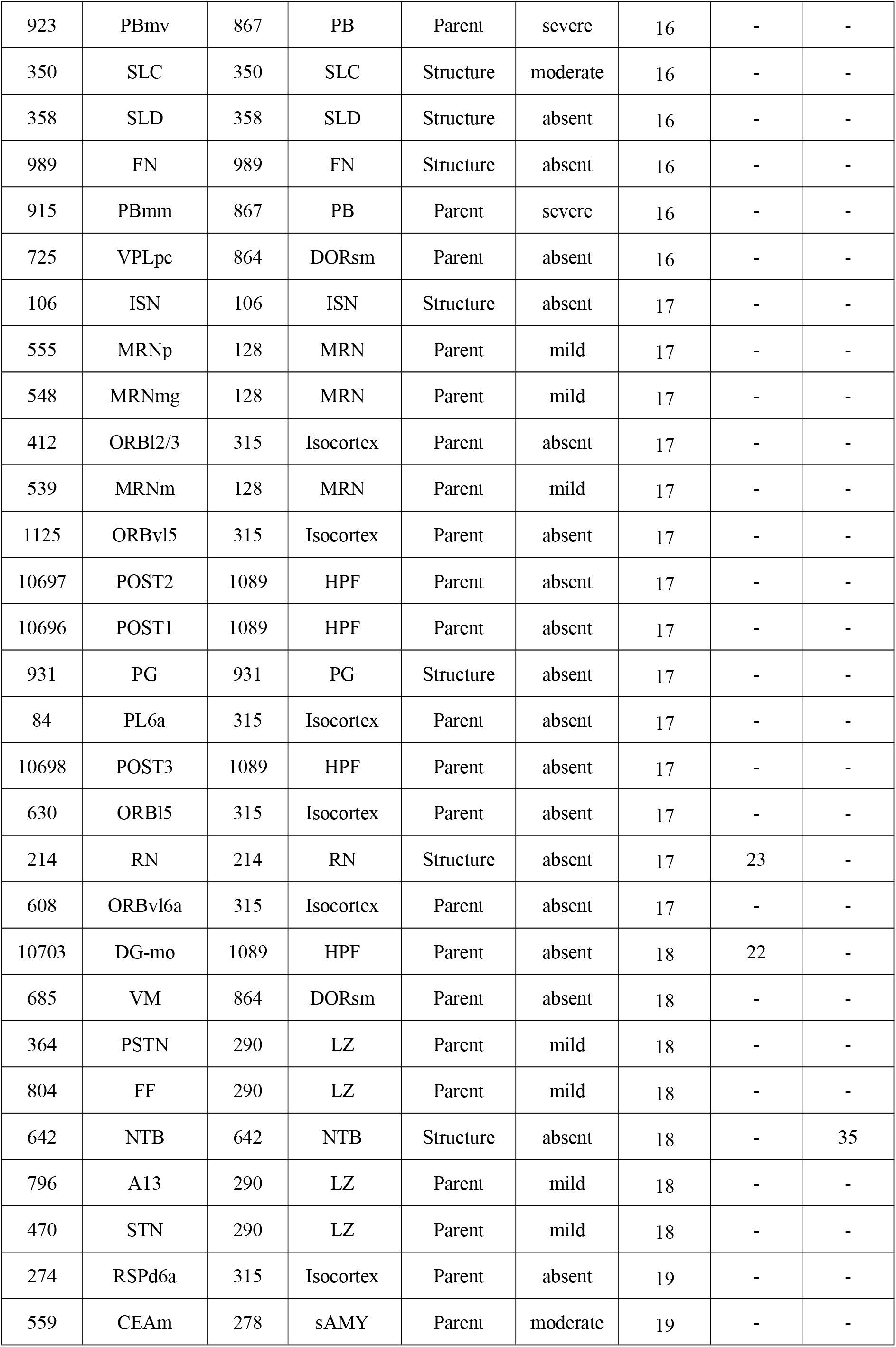
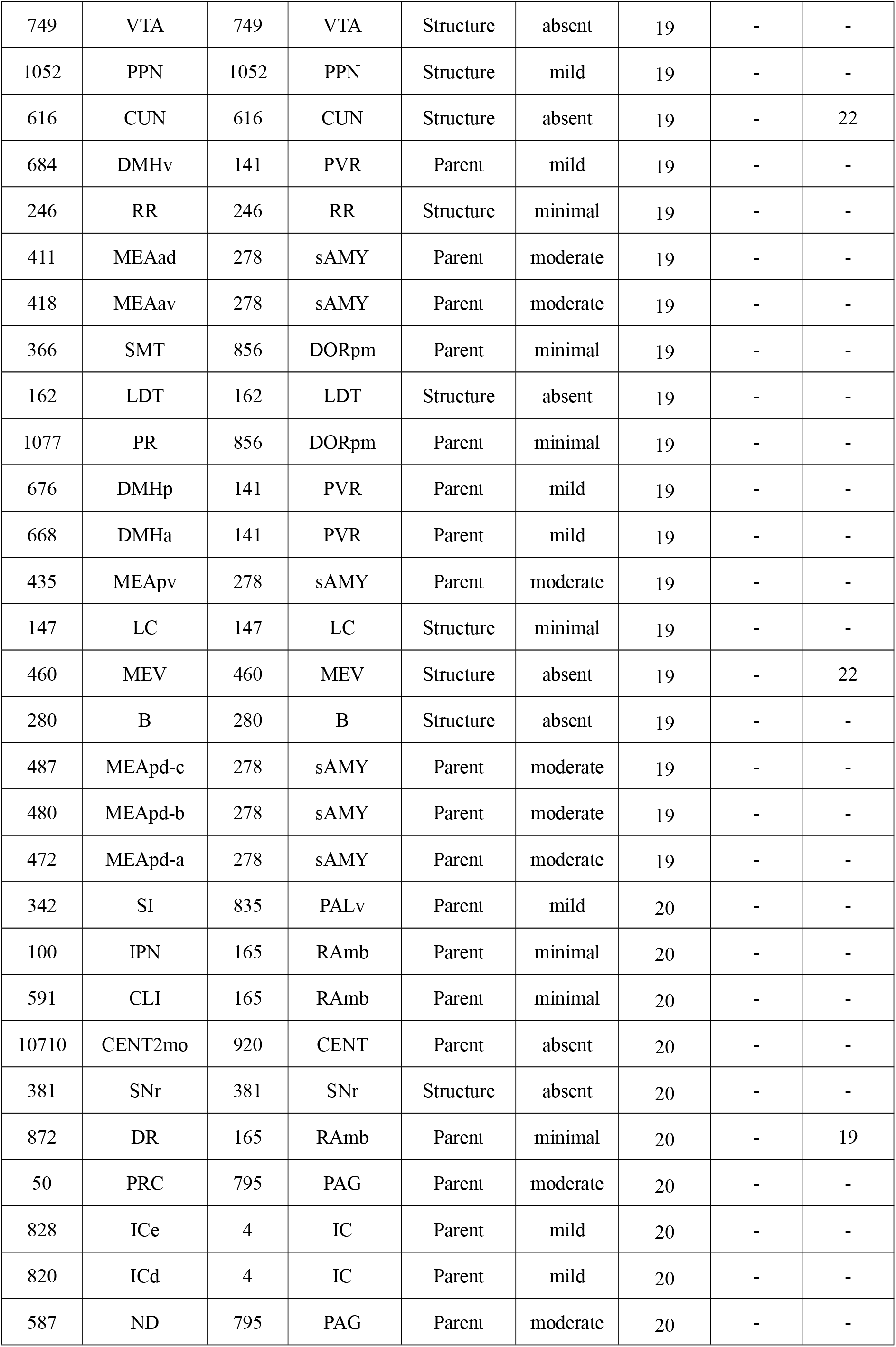
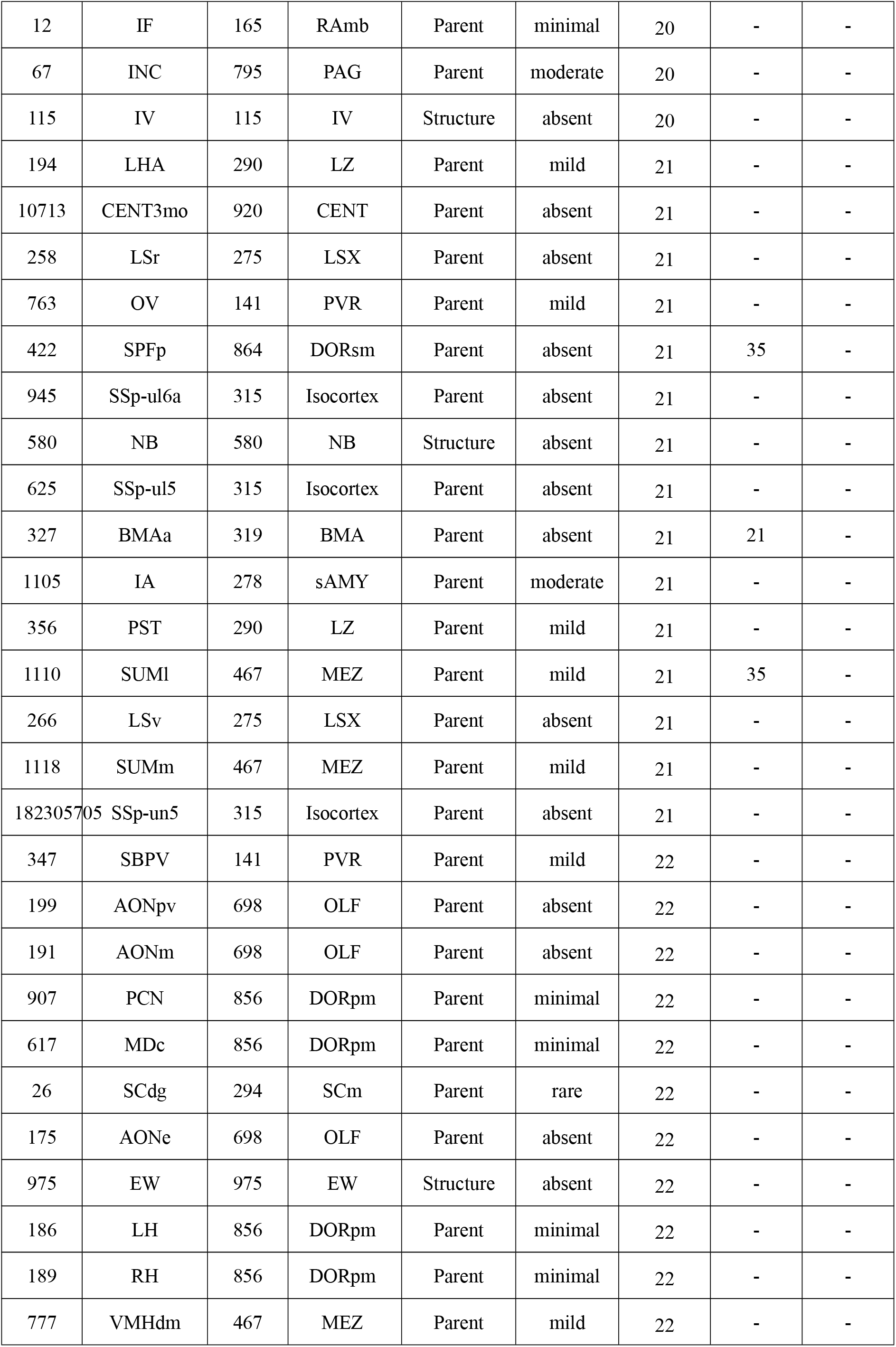
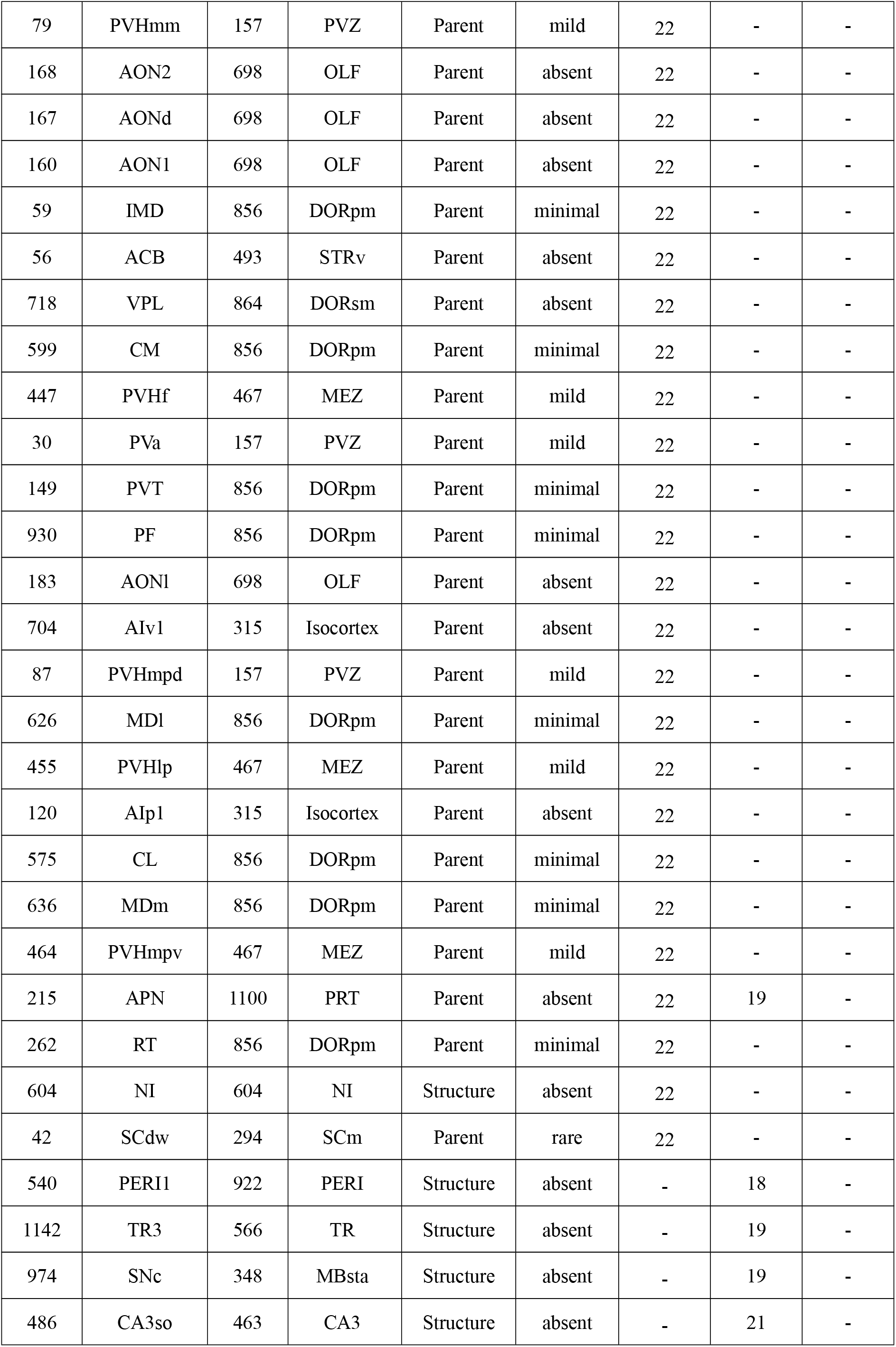
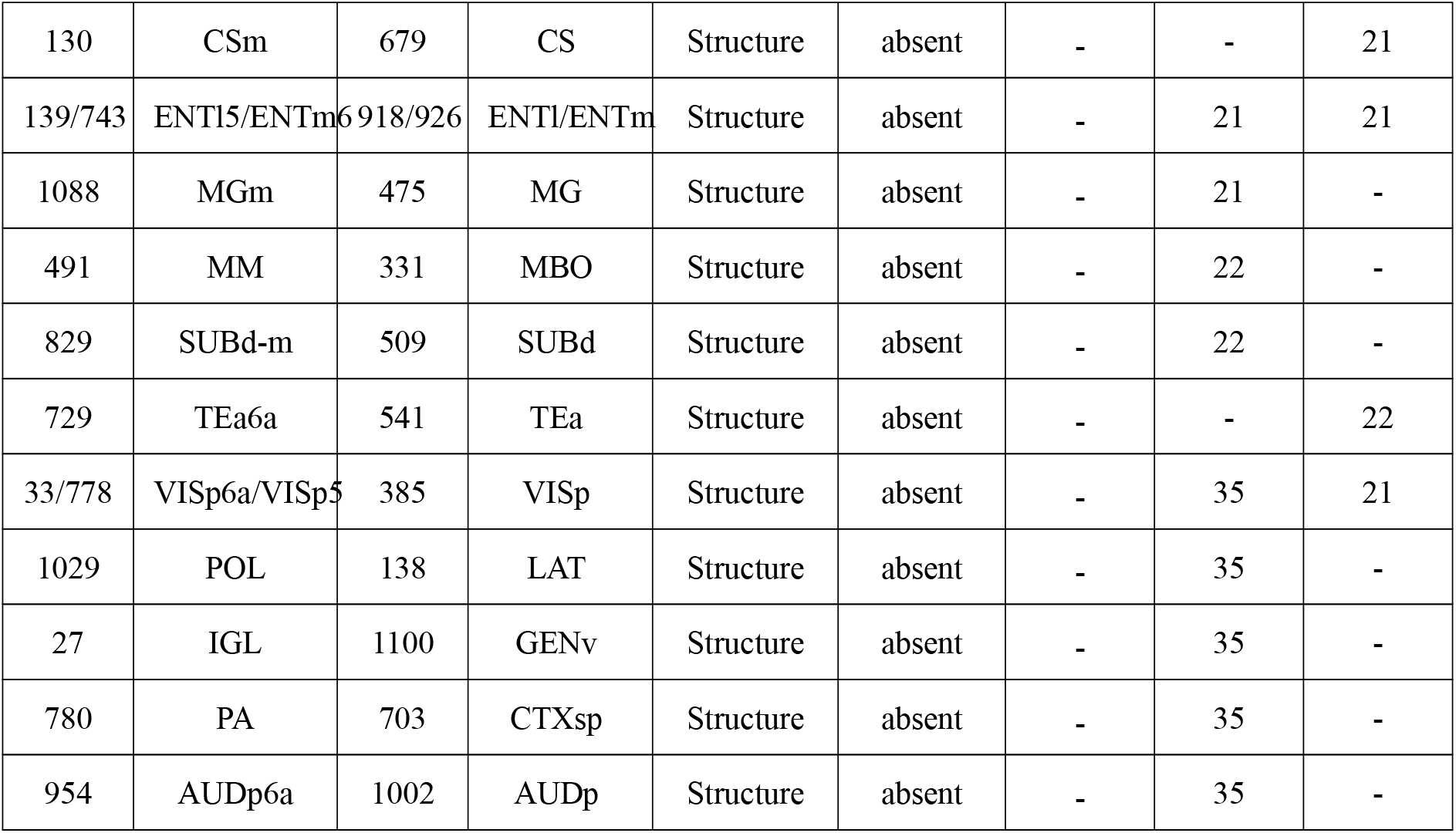
A comparison of *alpha*-synuclein spreading within a rat model (12 weeks after injection of *alpha*-synuclein expressing adeno-associated virus into the left vagus), and the first occurrence of virtual alpha-synuclein accumulation in a simulation of spread within the first-order composite mouse mesoscale connectome (with respect to DMX and AMB as initiation structures). The mouse simulation time-step is the simulation time where the simulated spread first reaches the corresponding brain structure. The single *Network spread* column is the result of a simulation that assumes a structure to be alpha-synuclein positive if virtual alpha-synuclein reached it via a source or target structure, ignoring the intersection of model projections with intermediate brain structures. In contrast, the two *coronal slice* columns are the results of simulations that assume a brain structure is alpha-synuclein-positive if virtual alpha-synuclein reaches it via a source, target, or intermediate structure along a model projection, only for those structures that are present the respective coronal slices.

### 3.5 Detailed comparison between whole-brain mouse simulation and synuclein pathology in Parkinson’s disease

**Table 3:**
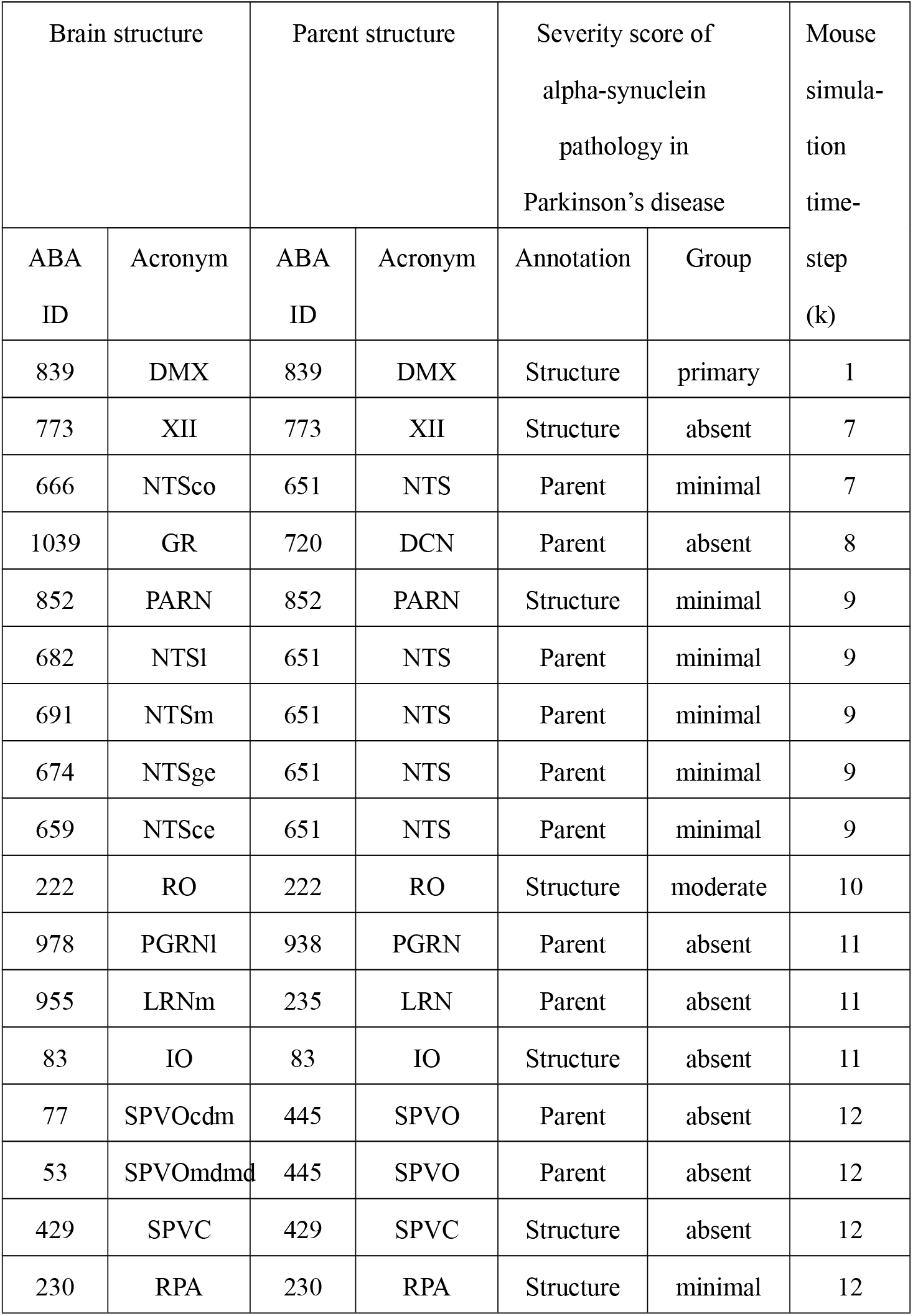
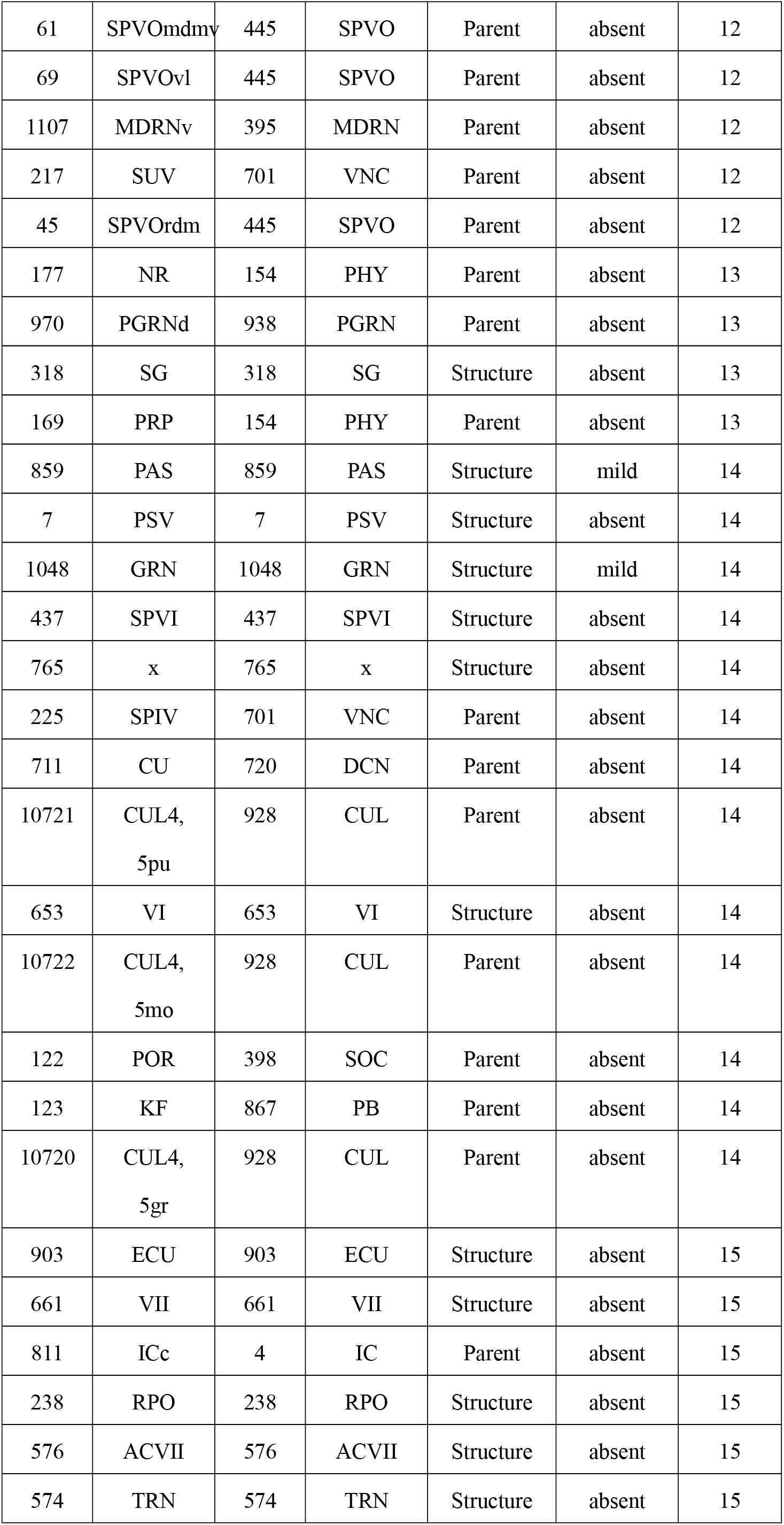
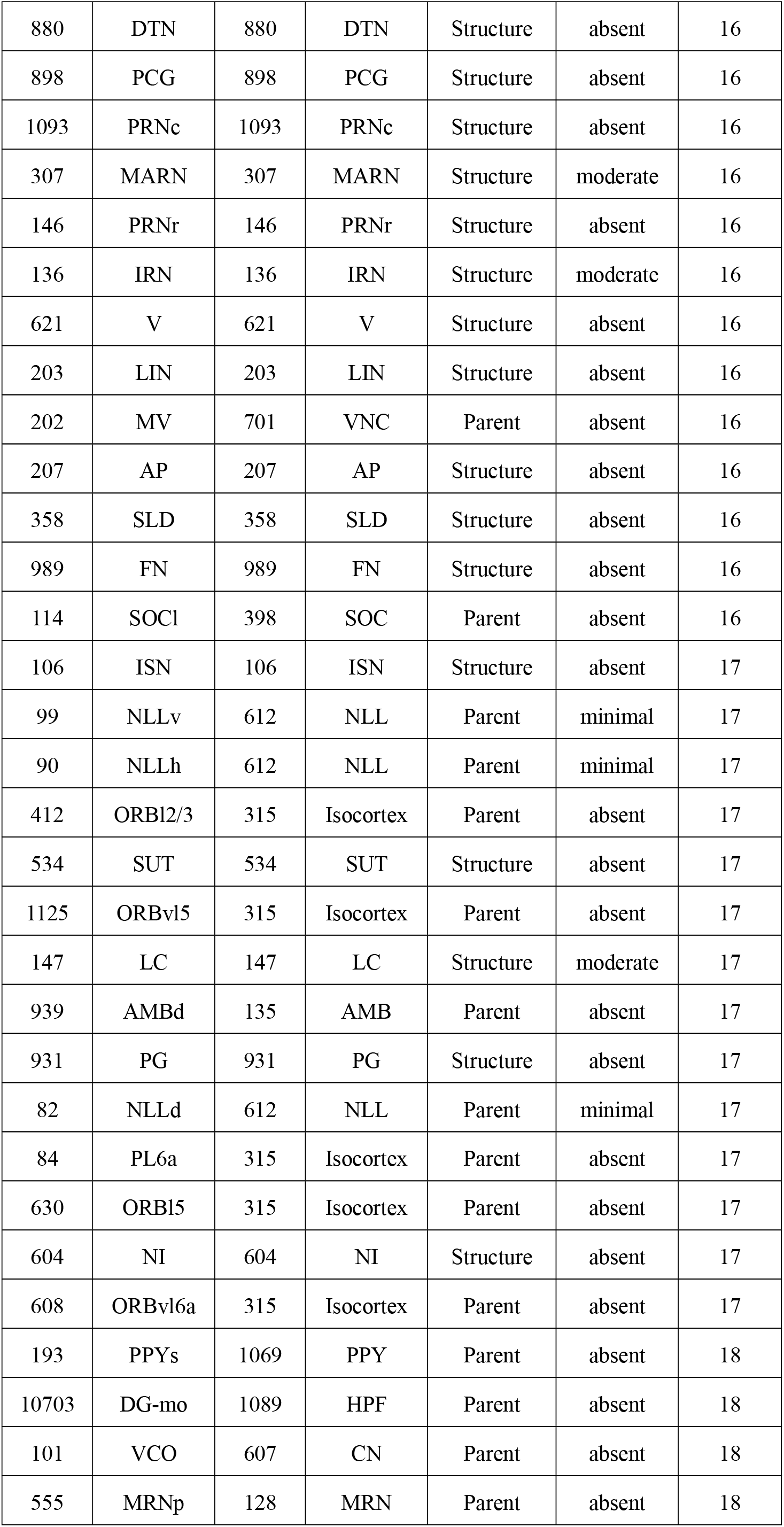
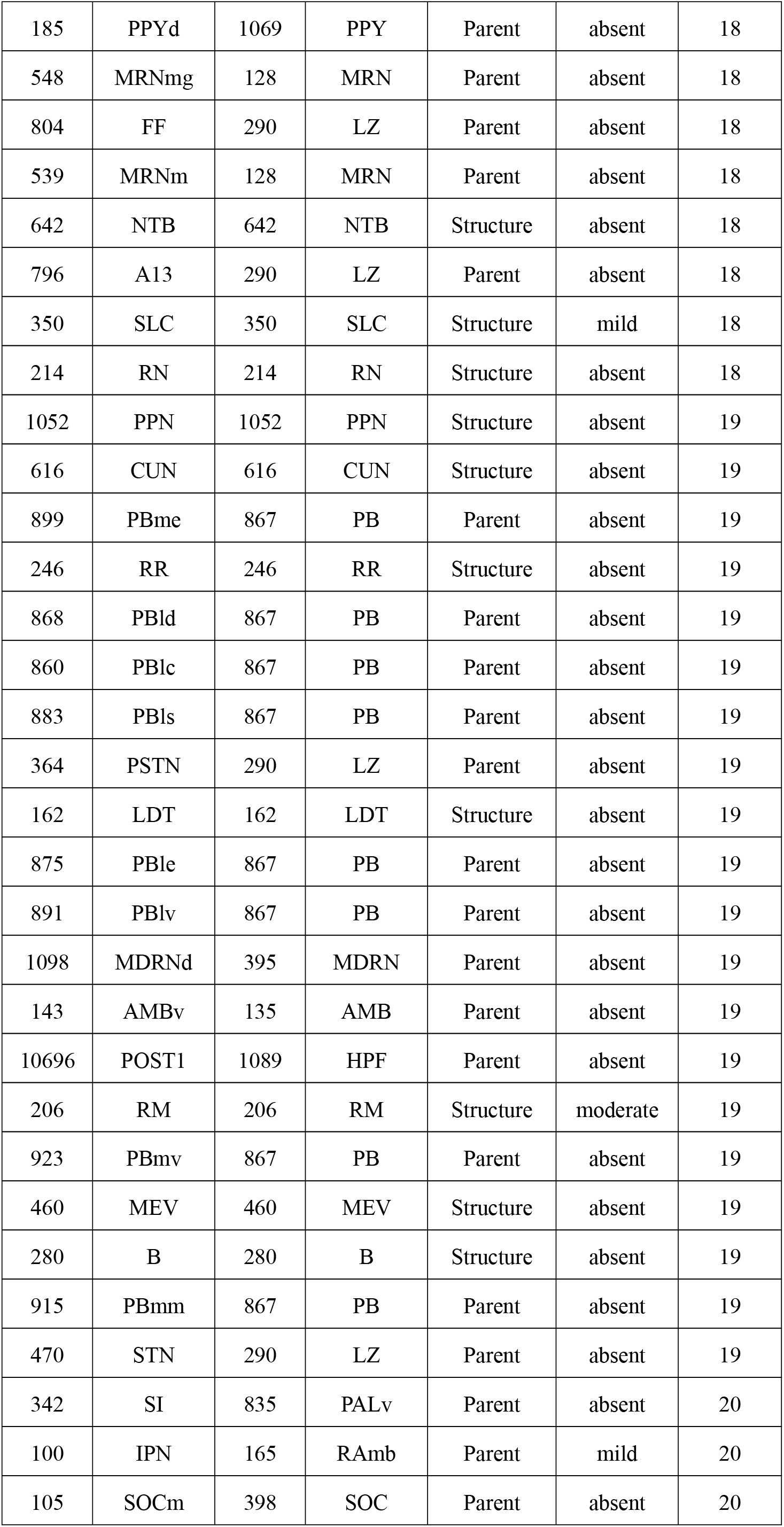
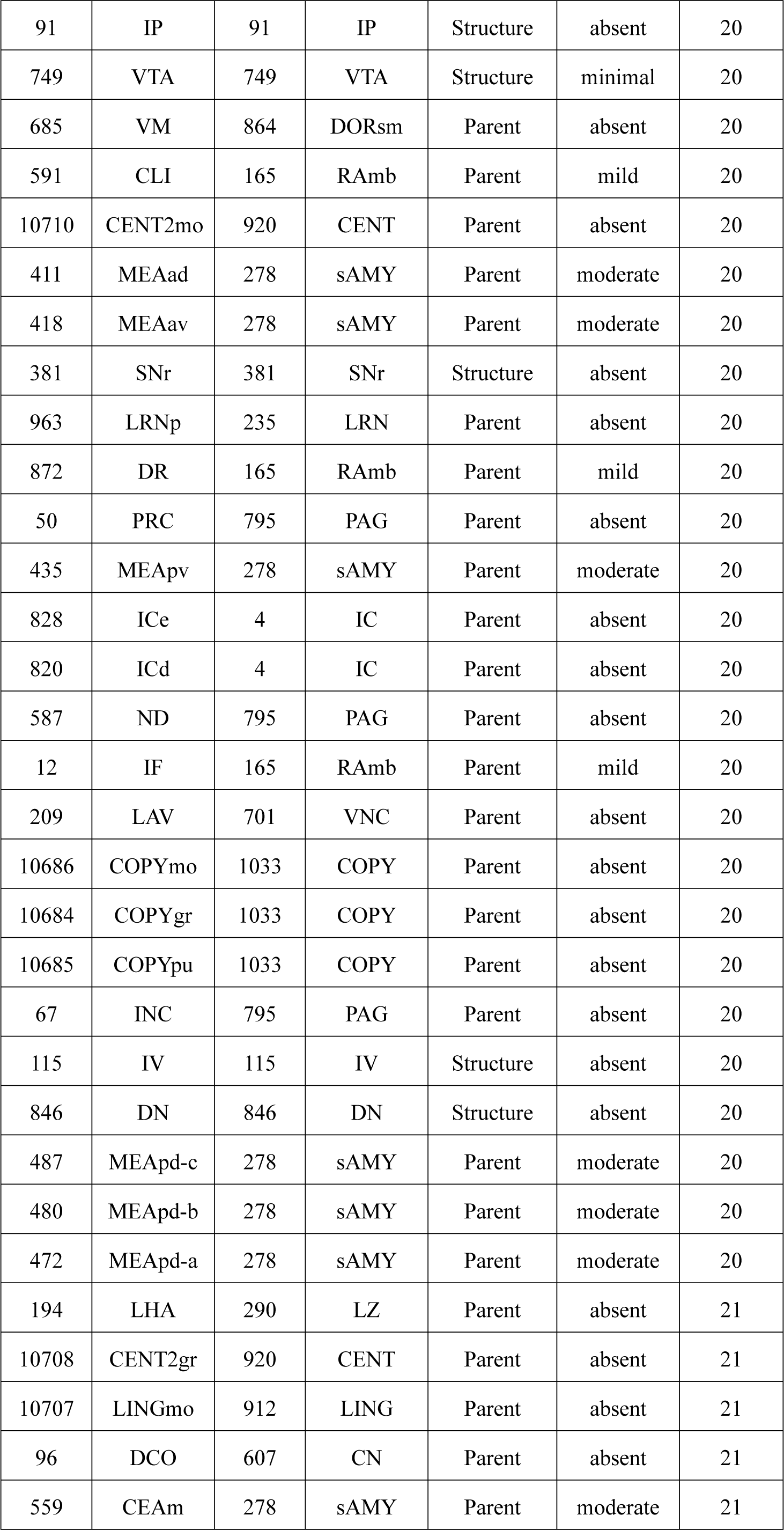
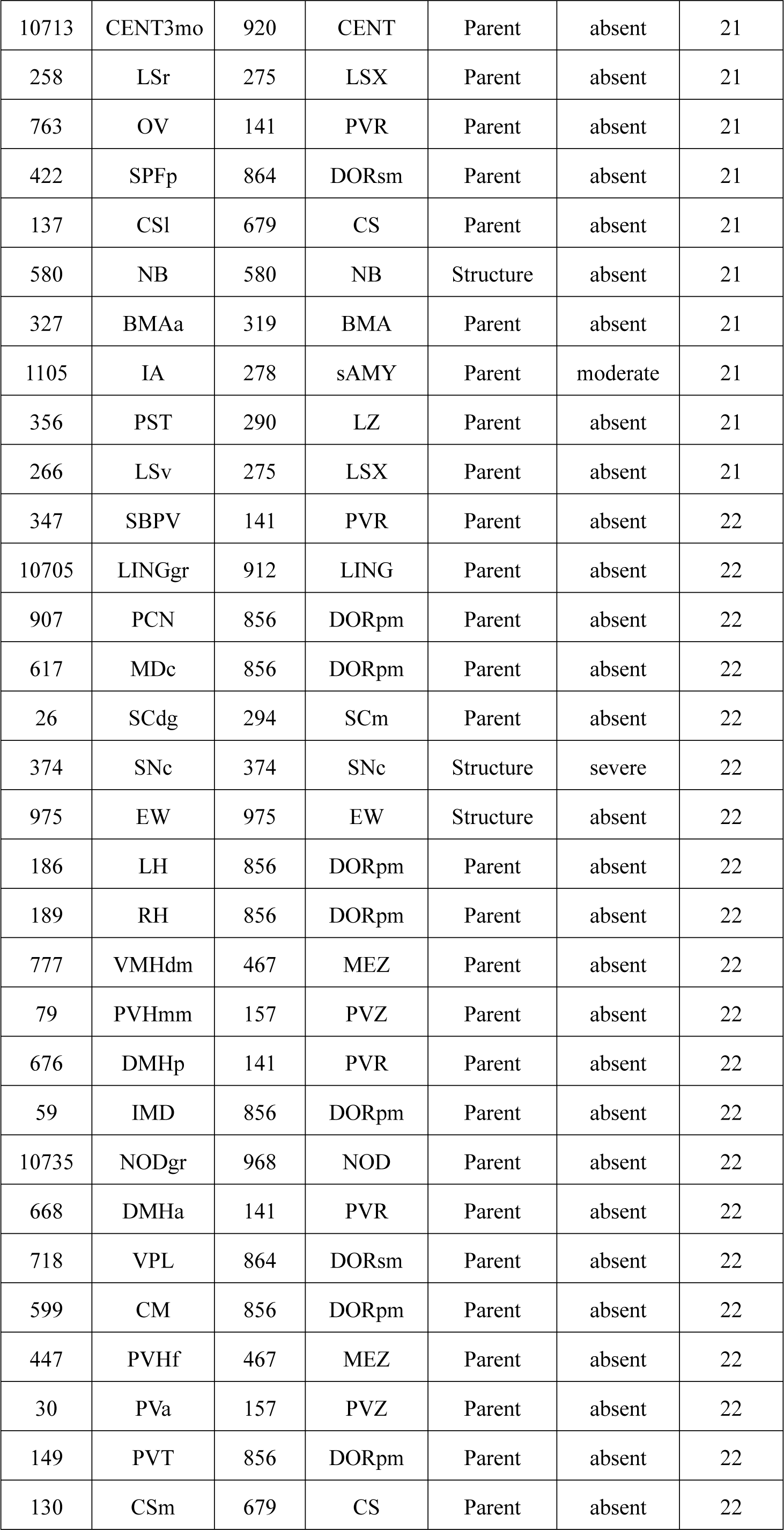
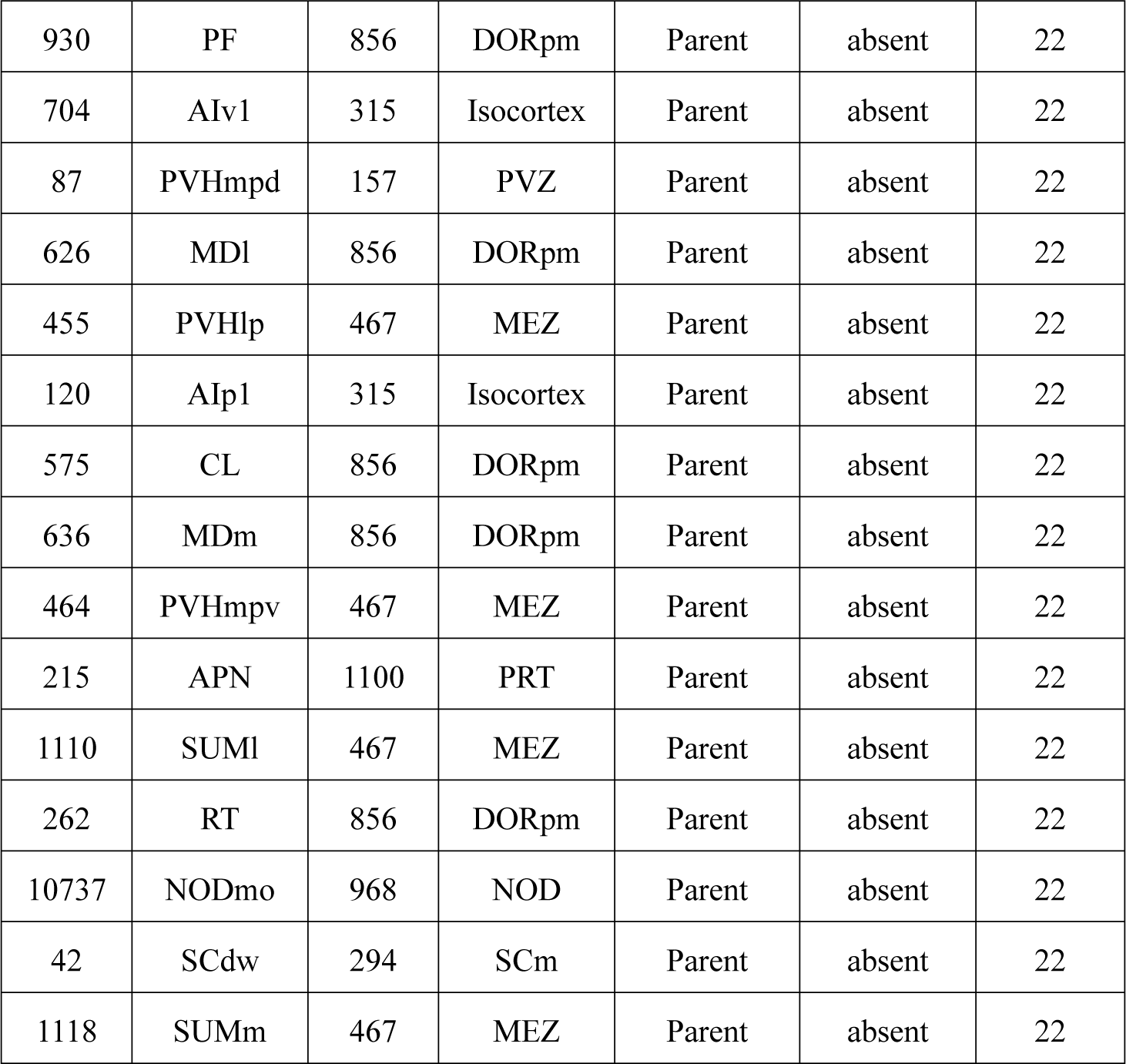
Detailed whole-brain simulated spread profile for the first 22 time-steps, and its correspondence with the annotated alpha-synuclein pathology in Parkinson’s disease (severe/ moderate/ mild/ minimal/ absent) (Surmeier et al. 2017; Oliveira et al. 2017), after injection of into the dorsal motor nucleus of the vagus. The mouse simulation time-step is the simulation time where the simulated spread first reaches the corresponding brain structure. A structure as *alpha*-synuclein positive if virtual *alpha*-synuclein reached it via a source or target structure, ignoring the intersection of model projections with intermediate brain structures.

## Supplementary Discussion

### 3.6 Protein aggregation and cell-to-cell transmission

Intracellular protein aggregates can be released from neurons by exocytosis or cell death. The aggregates are taken up by, for example, adjacent neuronal cell bodies and are either retained in the cell soma (local spread of pathology) or transported anterogradely by axons. Anterograde transmission along an axon is from the soma to the axonal terminal, and retrograde, the reverse. Alternatively, they are taken up by axon terminals and transported retrogradely to the cell soma. The protein can spread between brain regions by axonal transport. The spread of protein aggregates in the nervous system has been hypothesised to contribute to the advances of clinical symptoms and neuropathological changes in Alzheimer’s, Parkinson’s and Huntington’s diseases. Recent studies have demonstrated cell-to-cell or media-to-cell transfer of aggregates, but have given less insight into the mechanisms by which they are released or taken-up (Brundin et al. 2010).

Potential mechanisms for trans-cellular propagation of protein misfolding (Frost et al. 2010) are (a) intracellular protein aggregation resulting in release of proteinaceous seeds (aggregates or misfolded proteins), into the extracellular space, which are subsequently taken up and corrupt protein folding in vulnerable cells; (b) as part of the physiology of a living cell, proteinaceous seeds may be released, potentially via exosomes or exocytosis. This results in proteinaceous seeds in the extracellular space that may be taken up by adjacent cells. This mechanism can account for local propagation of misfolding. (c) Proteinaceous seeds might cross synapses. Release could be due to local degeneration of a synapse, normal synaptic physiology, or as part of an exocytotic process as in (b). This mechanism may explain network degeneration in neurodegenerative diseases.

